# Upstrap for Estimating Power and Sample Size in Complex Models

**DOI:** 10.1101/2021.08.21.457220

**Authors:** Marta Karas, Ciprian M. Crainiceanu

## Abstract

Power and sample size calculation are major components of statistical analyses. The upstrap resampling method introduced by Crainiceanu and Crainiceanu (2018) was proposed as a general solution to this problem but has not been assessed in numerical experiments. We evaluate the power estimation properties of the upstrap for target data sets that are larger or smaller than the original observed data set. We also expand the scope of upstrap and propose a solution to estimate the power to detect: (1) an effect size observed in the original data; and (2) an effect size chosen by a researcher. Simulations include the following scenarios: one- and two-sample t-tests; linear regression with both Gaussian and binary outcomes; multilevel mixed effects models with both Gaussian and binary outcomes. In addition, our simulations consider cases where the distribution of a covariate in the target setting is preserved and when it is purposefully changed compared to the original data set. We illustrate the approach using a reanalysis of a cluster-randomized controlled trial of malaria transmission. The GitHub repository with R code used in manuscript analyses is available at https://git.io/J0TH1. The accompanying data are publicly available.

## 1 Introduction

We consider the context when a test statistic is used on a set of preliminary data. We evaluate the power and sample size estimation properties of the test statistics using the upstrap (1), a general-purpose resampling technique. In summary, the upstrap starts with a sample (observed data) and re-samples with replacement either fewer or more samples than in the original data set. To be even more precise, here is the difference between bootstrap and upstrap.

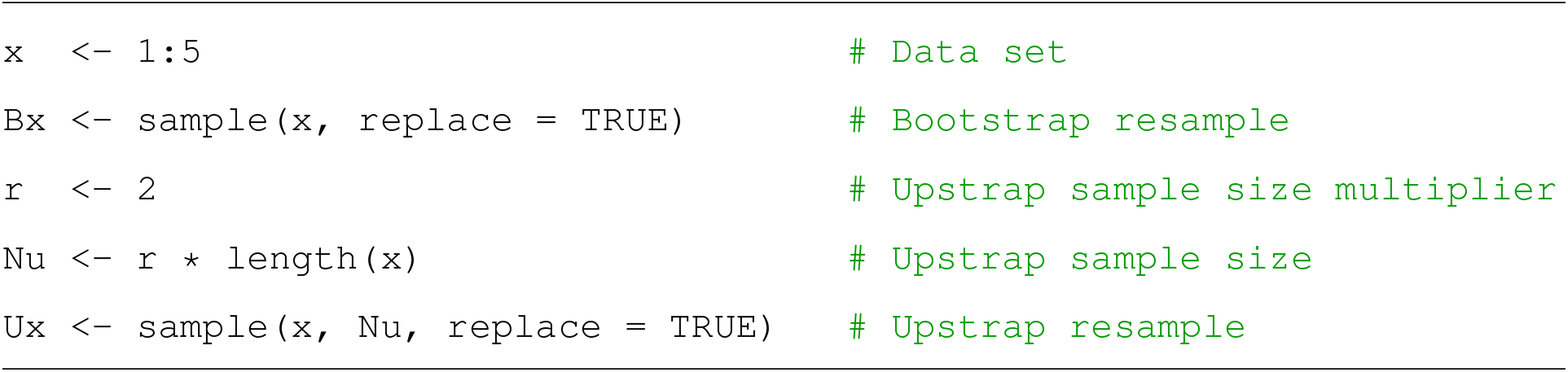

The estimator of a parameter of interest can be obtained from an upstrap resample for any model, procedure, or algorithm that provides estimators of a target parameter. A collection of these estimators can be obtained by generating multiple upstrap resamples. Such a collection provides a sample from the distribution of the estimator of a parameter of interest when the sample size is either larger or smaller than the original one.

We also expand the scope of the upstrap approach by proposing a solution to estimate the power to detect: (1) an effect size observed in the preliminary data; and (2) effect size chosen by a user. We present “ready to use” code examples to estimate power and sample size to identify a statistically significant effect for an α-level test to achieve the power 1 – *β*. We evaluate the accuracy of the upstrap to estimate power in an extensive simulation study. We consider both the case when analytic solutions are available (e.g., one-sample t-test) and when they are not (e.g., testing for fixed effect in a multilevel mixed-effects model). In addition, our simulations consider cases where the distribution of a dichotomous covariate in the target setting is preserved and when it is purposefully changed compared to the original data set. We illustrate the approach in a reanalysis of results from a cluster-randomized controlled trial of the impact of interventions on malaria transmission (2), where the treatment was assigned at the cluster level.

### 1.1 Related work

#### 1.1.1 Nonparametric boostrap for multilevel data

The nonparametric bootstrap for multilevel data is one of the accepted analytic methods for quantifying variability in mixed effects modeling. For example, (3) considered hierarchical data with two levels; they proposed to first randomly sample groups with replacement and then to sample within the groups without replacement (“case bootstrap”). (4) extended the concept to hierarchical unbalanced data with two and more levels. They showed theoretically that sampling units with replacement at the highest level and then sampling within these units without replacement resembles the variation properties of the data more closely than sampling units at lower hierarchical levels. They also showed in simulations (two- and three-level data cases) that sampling units at the highest level gave more accurate estimates for the highest level fixed-effect parameter. (5) used simulations to evaluate a range of bootstrap methods for estimating confidence intervals and standard errors of parameters in a linear mixed-effect model (two-level balanced data case). They provided evidence that overall, the case bootstrap outperforms or performs similarly to other approaches they considered (case bootstrap coupled with global/individual residual bootstrap, random effects bootstrap coupled with global/individual residual bootstrap, global/individual residual bootstrap and parametric bootstrap). (6) used the case bootstrap of two-level unbalanced data for standard error estimation in a context that mimics a cluster-randomized controlled trial. They proposed a modification of separating the “treatment” and “control” clusters into two parts within which step 1 of the case bootstrap is done independently.

#### 1.1.2 Power analysis and sample size estimation for multilevel data

The power analysis and sample size calculations for multilevel data are active areas of research. *A priori* power analysis aims at estimating the sample size needed to achieve specific power 1 – *β* for a given significance level *α* and hypothesized effect size. *Post hoc* power analysis is conducted using available data to estimate the power of a test and the sample size needed for a future study (7).

A class of theoretical results for estimating power in specific multilevel data setups has been proposed; see, for example, references in (8). These results are based on the variance inflation factor correction approach (9; 10). The approach uses assumptions about the intra-class correlation coefficient, *ρ*, or assumes a particular study design (e.g., that the data are balanced).

Another class of approaches is based on simulations. (7) proposed a general approach that requires specifying a population model for the data, simulating data from the assumed model, and estimating the power via Monte Carlo simulations. There exists software designed for *post-hoc* power analysis of a test given a multilevel data and model (11). For example, the `SIMR` R package (12; 13) is a versatile R software for power analysis of generalized linear mixed models (GLMMs) using simulation. It allows estimating a power curve for a range of sample sizes by simulating new values for the response variable and refitting the model to the larger data set.

To the best of our knowledge, this is the first time upstrap resampling techniques have been proposed for power analysis and sample size estimation in GLMMs. The remainder of the paper is organized as follows. Section 2 describes the upstrap approach for estimating power and sample size. Section 3 presents the simulation study. Section 4 describes the applications of our approach to the reanalysis of cluster-randomized controlled trial data on the impact of hotspot-targeted interventions on malaria transmission (2). We close with a discussion in Section 5. Supplementary Material provides “read, adapt and use” R code for estimating power with upstrap in a series of examples, and also contains a part of simulation results. The GitHub repository with R code for all manuscript analyses is available at https://git.io/J0TH1.

## 2 Methods

### 2.1 Notation

Denote by **x** a vector of *N* independent realizations of a random variable *X* with an unknown distribution *F*. We assume that **x** consists of either scalar-valued, *x_i_*, or vector-valued, **x***_i_*, observations. Observations are assumed to be independent across observational units, *i* = 1, …, *N*. Throughout the manuscript, the term “sample size” denotes the number of independent units in **x**. When observations are scalar and independent, the sample size is the number of observations. When observations are vectors with a multilevel dependence structure, the sample size denotes the number of unique units at the highest level of the data structure.

### 2.2 The upstrap

The upstrap method (1) samples with replacement from **x** either more or fewer samples than the original sample size, *N*. For a fraction *r* ∈ (0, ∞), the upstrap creates *B* samples, 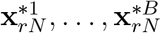, by sampling with replacement from **x** with a sample size *rN*. The estimators 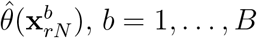 can be obtained for any model, procedure, or algorithm that provides estimators of a target parameter *θ*. The collection of these estimators provides a sample from the distribution of the estimator 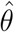 for a dataset of size *r* times the sample size *N* of the original data.

### 2.3 The upstrap solution to the sample size estimation problem

We consider the following problem: given **x** of sample size *N* from unknown distribution *F*, a specific null and alternative hypothesis, a test statistic, and significance level *α* of the test, estimate a sample size *M* required to achieve probability 1 – *β* of rejecting the null hypothesis when the null is true (i.e., achieve power 1 – *β*).

#### 2.3.1 Target effect size as observed in preliminary data x

In case the target effect size is the one observed in sample **x**, we propose the following solution:

**Step 1:** Consider a grid of target sample sizes, *M_k_*, for *k* = 1, …, *K*, where *K* is an integer denoting the grid length.
**Step 2:** Use the upstrap method to estimate the power for a sample of size *M_k_*, for *k* = 1, …, *K*:

a. Generate *B* resamples of size *M_k_*, denote 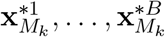, by sampling with replacement from **x**;
b. Perform hypothesis test on each resample 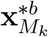, for *b* = 1, …, *B*;
c. Define 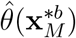 to be equal to 1 if the null hypothesis was rejected for data 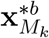 and equal to 0 otherwise, for *b* = 1, …, *B*;
d. Define 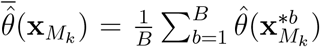 the proportion of resamples when the null hypothesis was rejected as the estimate of power.
**Step 3:** Identify the smallest sample size *M_k_* for which the desired power 1 – *β* is achieved.

Denote by 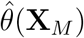 the sampling distribution of the statistic 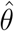 when derived from a random sample **X** of size *M*. For a given realization **x** of sample size *N*, a sample mean of the upstrap-generated values 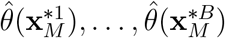, denote 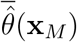, is an estimate of the mean of 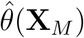 distribution and the proposed estimate of power.

#### 2.3.2 Arbitrary target effect size

A common practical case is when the target effect size is larger or smaller than the one observed in the preliminary data. To address such cases, we propose the following approach:

**Step 0:** Update the observed outcome values to ensure that the effect size in this updated data is the [target effect size × target covariate value] instead of the [observed effect size × target covariate value].

For general linear model (GLM)/GLMM with an identity link function, Step 0 is fairly straightforward to implement. Specifically, we propose to update each observation in sample **x** by adding to the outcome the value of [(target effect size – observed effect size) × target covariate value]. The observed effect size is computed once for **x** at the beginning of the procedure, and then the updated sample **x** is used in Steps 1-3. See R code examples in Supplementary Material in Sections 1.1, 1.2, 1.3, 1.5.

For other distributions in the GLM/GLMM family (e.g., Gamma, Poisson or Bernoulli), we propose the following modification. For each upstrap resample from Step 2a: (1) use the resample to fit the assumed model; (2) update the obtained model fit by setting the coefficient of interest to [target effect size]; (3) use the updated fit to simulate new outcome values for the resample; and (4) use the updated resample in Step 2b. See R code examples in Supplementary Material in Section 1.6. The idea of simulating a new outcome based on an updated R model fit object is shared with the `SIMR` approach. For some samples with independent observations, the procedure can be simplified by simulating new outcome values from the distribution instead of R model fit. Here, the distribution parameters are based on the linear predictor estimated for original sample after updating the coefficient of interest to [target effect size]. See R code examples in Supplementary Material in Section 1.4.

#### 2.3.3 Sampling with replacement from x

When the observed sample **x** consists of independent scalar observations, the sampling with replacement from **x** is done by sampling the observations. When observations are vectors with a multilevel dependence structure we recommend the “case sampling”, i.e. sample units at the highest level with replacement, and then sample within these units without replacement at the lower level.

## 3 Simulations

An extensive simulation study was used to assess the performance of the upstrap to estimate the power of an α-level test for a sample size smaller or larger than in an observed sample. Overall, simulations span three different contexts for evaluating power estimation: (a) detecting both arbitrary effect size and effect size observed in the data in simple and complex models, and comparing upstrap with existing methods (problems 1-6); (b) scenarios where the target distribution of a dichotomous covariate is changed compared to the original data set (problems 7-9); (c) scenarios with a very small observed sample size to evaluate limits of upstrap good performance (problems 10-12). The R code for the simulation study is provided on the project’s GitHub repository (sub-directory URL: https://git.io/JsiiA).

### 3.1 Setup of the problems

Table 1 summarizes the setup for 12 problems considered in the simulations. The detailed specification of data-generating models is included in Table A.2 in Appendix A. In all simulations, a two-sided test was used to test *H*_0_ : *β* = 0 versus *H*_1_ : *β* ≠ 0 at significance level *α* = 0.05.

**Table 1:**
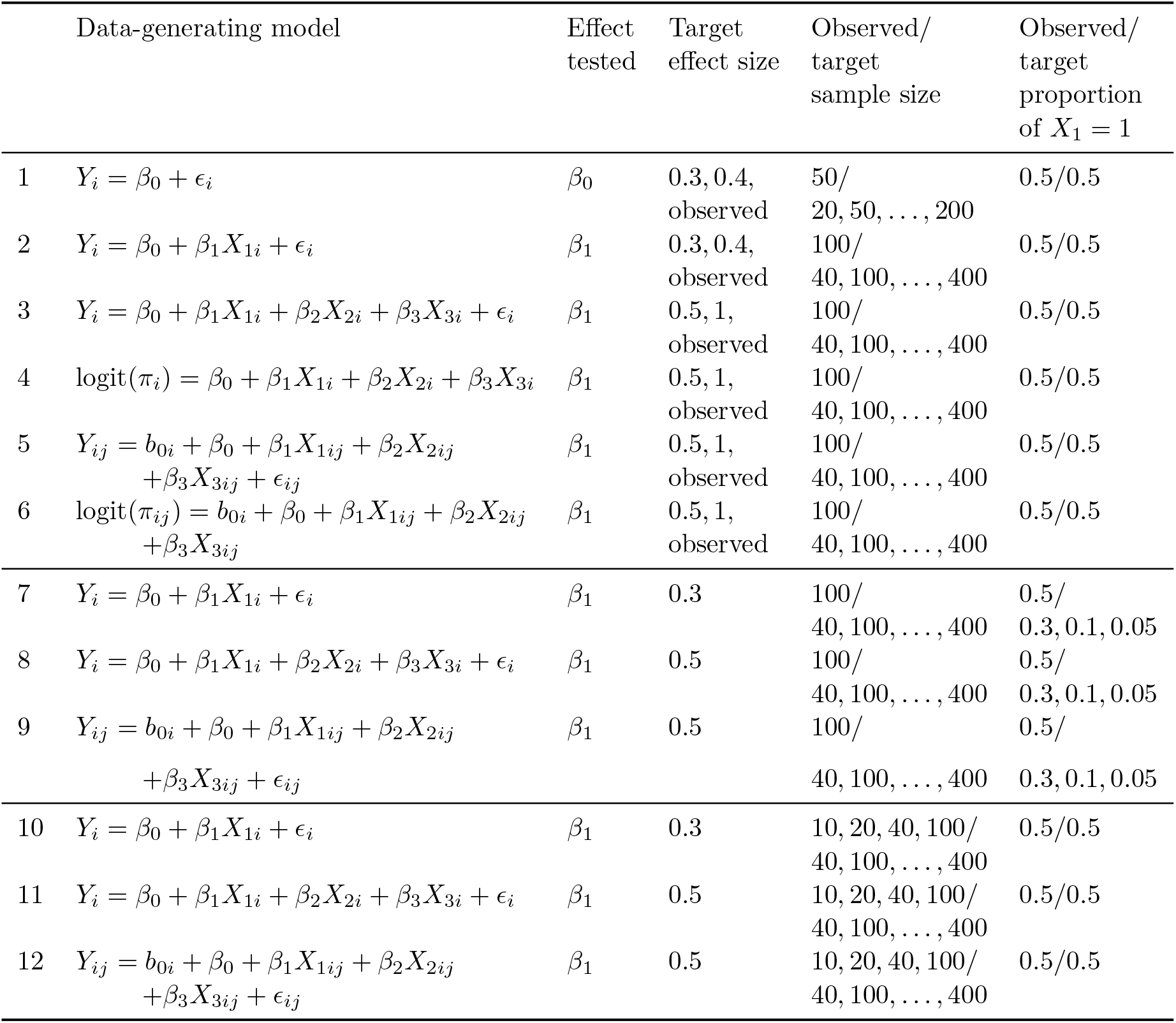
Summary of simulation setup across 12 different problems. In all cases, “sample size” refers to the number of independent units in the sample indexed by *i*.

| | Data-generating model | Effect tested | Target effect size | Observed/<br>target<br>sample size | Observed/<br>target<br>proportion<br>of $X_1 = 1$ |
| --- | --- | --- | --- | --- | --- |
| 1 | $Y_i = \beta_0 + \epsilon_i$ | $\beta_0$ | 0.3, 0.4,<br>observed | 50/<br>20, 50, ..., 200 | 0.5/0.5 |
| 2 | $Y_i = \beta_0 + \beta_1 X_{1i} + \epsilon_i$ | $\beta_1$ | 0.3, 0.4,<br>observed | 100/<br>40, 100, ..., 400 | 0.5/0.5 |
| 3 | $Y_i = \beta_0 + \beta_1 X_{1i} + \beta_2 X_{2i} + \beta_3 X_{3i} + \epsilon_i$ | $\beta_1$ | 0.5, 1,<br>observed | 100/<br>40, 100, ..., 400 | 0.5/0.5 |
| 4 | $\text{logit}(\pi_i) = \beta_0 + \beta_1 X_{1i} + \beta_2 X_{2i} + \beta_3 X_{3i}$ | $\beta_1$ | 0.5, 1,<br>observed | 100/<br>40, 100, ..., 400 | 0.5/0.5 |
| 5 | $Y_{ij} = b_{0i} + \beta_0 + \beta_1 X_{1ij} + \beta_2 X_{2ij} + \beta_3 X_{3ij} + \epsilon_{ij}$ | $\beta_1$ | 0.5, 1,<br>observed | 100/<br>40, 100, ..., 400 | 0.5/0.5 |
| 6 | $\text{logit}(\pi_{ij}) = b_{0i} + \beta_0 + \beta_1 X_{1ij} + \beta_2 X_{2ij} + \beta_3 X_{3ij}$ | $\beta_1$ | 0.5, 1,<br>observed | 100/<br>40, 100, ..., 400 | 0.5/0.5 |
| 7 | $Y_i = \beta_0 + \beta_1 X_{1i} + \epsilon_i$ | $\beta_1$ | 0.3 | 100/<br>40, 100, ..., 400 | 0.5/<br>0.3, 0.1, 0.05 |
| 8 | $Y_i = \beta_0 + \beta_1 X_{1i} + \beta_2 X_{2i} + \beta_3 X_{3i} + \epsilon_i$ | $\beta_1$ | 0.5 | 100/<br>40, 100, ..., 400 | 0.5/<br>0.3, 0.1, 0.05 |
| 9 | $Y_{ij} = b_{0i} + \beta_0 + \beta_1 X_{1ij} + \beta_2 X_{2ij}$ | $\beta_1$ | 0.5 | 100/ | 0.5/ |

| | $+\beta_3 X_{3ij} + \epsilon_{ij}$ | | | 40, 100, \dots, 400 | 0.3, 0.1, 0.05 |
| --- | --- | --- | --- | --- | --- |
| 10 | $Y_i = \beta_0 + \beta_1 X_{1i} + \epsilon_i$ | $\beta_1$ | 0.3 | 10, 20, 40, 100/<br>40, 100, \dots, 400 | 0.5/0.5 |
| 11 | $Y_i = \beta_0 + \beta_1 X_{1i} + \beta_2 X_{2i} + \beta_3 X_{3i} + \epsilon_i$ | $\beta_1$ | 0.5 | 10, 20, 40, 100/<br>40, 100, \dots, 400 | 0.5/0.5 |
| 12 | $Y_{ij} = b_{0i} + \beta_0 + \beta_1 X_{1ij} + \beta_2 X_{2ij}$<br>$+\beta_3 X_{3ij} + \epsilon_{ij}$ | $\beta_1$ | 0.5 | 10, 20, 40, 100/<br>40, 100, \dots, 400 | 0.5/0.5 |

### 3.2 Comparator methods

For simulation problems 1-6, the upstrap approach for power estimation was compared with an existing method.

#### 3.2.1 Simulation problems 1-2

Hypothesis testing in simulation problems 1-2 narrows down to one-sample and two-sample t-test, respectively, for which analytic solutions for power estimation exit. Their implementation, available in `power.t.test()` R function, was used as a comparator approach. Specifically, for the case when the target effect size was set to the effect size observed in the sample **x**, sample statistics computed from an observed sample **x** (mean/difference of means, standard deviation) were used as `power.t.test()` function’s arguments `delta` and `sd`. For the case when the target effect size was set to an arbitrary value, that value was used as `delta` argument.

#### 3.2.2 Simulation problems 3-6

For simulation problems 3-6, the `SIMR` R package (12; 13) was used as a comparator. As discussed in (12), “in `SIMR`, power is calculated by repeating the following three steps: (i) simulate new values for the response variable using the model provided; (ii) refit the model to the simulated response; (iii) apply a statistical test to the simulated fit.”

To estimate power for sample size larger than that of the observed sample, the `extend()` function adds (upsample) rows to an original model data frame, creating the frame’s “extended” version. Then, `powerCurve()` estimates power at a range of sample sizes smaller than the “extended” data frame, including the sizes smaller than in the original data frame. To the best of our understanding, adding the rows in `extend()` is done once for the whole subsequent power estimation procedure. Also, in the `powerCurve()` function, the target sample size is achieved by removing rows from the “extended” data frame deterministically, i.e. by keeping the first (in row number order) observations. Therefore, the same set of observations will be used in the `SIMR` power estimation procedure across all iterations when simulating new values of the response. This is different from the upstrap approach, where data observations are resampled with replacement multiple times within the procedure.

Furthermore, for cases when the target effect size is the effect size observed in the sample **x**, `SIMR` uses a model fitted to **x** to simulate new responses. For cases when the target effect size is set to an arbitrary value, `SIMR` updates a model fitted to **x** by modifying its parameters according to the target effect value, and uses this updated model to simulate new responses. Finally, “`SIMR` is designed to work with any linear mixed model (LMM) or GLMM that can be fit with either `lmer` or `glmer` from `lme4`.”

To ensure a fair comparison between `SIMR` and upstrap several precautions have been used. First, both the upstrap and `SIMR` used the same `lmer` function for model coefficient testing in simulation problems 3-6. Second, the number of upstrap resamples (upstrap method’s parameter) was set to be equal to the number of simulations in `SIMR` (`SIMR` method’s parameter). Third, since there is no feature to maintain balance of covariates explicitly with the `SIMR powerCurve()` function, data samples were simulated to ensure that the balance of the dichotomous covariate of interest is preserved when `SIMR` removes samples. Fourth, the authors of the `SIMR` package and the (12) paper were contacted to ensure that `SIMR` is correctly used. One of the authors reported that “has not seen any obvious problems” with how we employed `SIMR`.

### 3.3 True power estimation

For all simulation problems, true power estimates were obtained by setting the generative model’s coefficient equal to the target effect size and repeatedly (10,000 times) sampling independent observations from the target sample size and the target covariates distribution. True power estimate was defined as the proportion of repetitions where null hypothesis was rejected and was denoted as 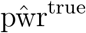.

### 3.4 Simulation procedure

The following procedure was performed to estimate and compare power at a range of target sample sizes for upstrap and a comparator approach, for each of the 12 simulation problems separately.

1. Consider *R* independent experiment repetitions. For each *r*-th repetition, *r* = 1, …, *R*, do:

a. Simulate an observed sample size **x***_r_*.
b. Use the upstrap to estimate power at a range of target sample sizes, 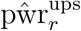.
c. Use the comparator approach to estimate power for a range of target sample sizes, 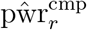,
d. Compute percentage error between:

i. upstrap and true power, 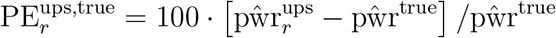;
ii. comparator and true power, 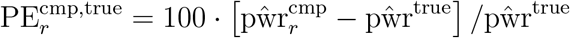;
iii. upstrap and comparator, 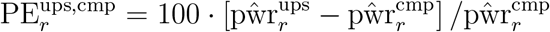. The *r*-th experiment repetition estimates: 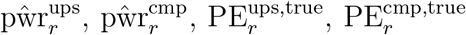 and 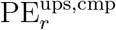 are vectors of the length equal to the number of target sample sizes considered.
2. Aggregate estimates from *R* independent experiment repetitions by computing mean, standard deviation, median, 25th percentile and 75th percentile.

Steps (1)(c) and (1)(d)(ii)-(iii) were done for simulation problems 1-6 only where a comparator approach was considered. The number of *R* independent experiment repetitions was set to 1000 for each simulation problem. The number of upstrap resamples (upstrap method’s parameter) and the number of simulations in `SIMR` (`SIMR` method’s parameter) were both set to 1000 in each case.

### 3.5 Simulation results

#### 3.5.1 Simulation problems 1 and 2

Figure 1 displays the power trajectories obtained with the upstrap (left plot) and comparator `power.t.test()` (right plot) in the first 15 experiment repetitions in simulation problem 1. The case when the target effect size is the effect size observed in the sample is considered. Repetition identifiers are indicated as red numbers. Figure 1 demonstrates that upstrap power estimates are very similar to those obtained from the `power.t.test()`. The difference is that they are slightly noisier, but this could be addressed by increasing the number of upstrap resamples.

**Figure 1:**
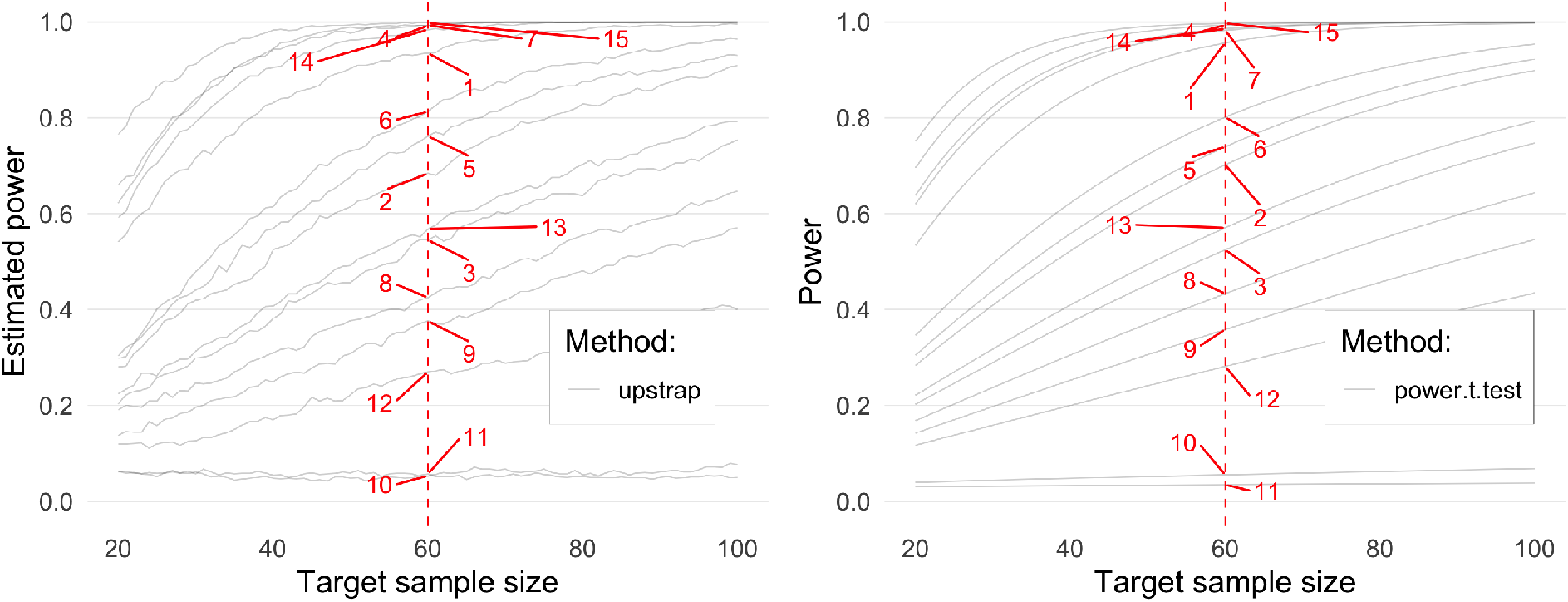
Power estimation for 15 repetitions from the simulation experiment 1. Left panel: upstrap, right panel: `power.t.test()` comparator approach. Red numbers denote a specific experiment repetition. Red lines point at power estimates for a target sample size equal 60 (chosen arbitrarily for presentation purposes).

Table SM.1 in Supplementary Material shows a summary of power estimates and percentage errors (PEs) across simulation problems 1-6. Figure 2 shows power estimates at a range of target sample sizes in simulation problem 1 (one-sample t-test) and problem 2 (two-sample t-test). In each case, the orange and purple points and lines show very close agreement, indicating that the results obtained based on the upstrap are essentially identical to those obtained based on a well-established analytical solution from `power.t.test()`. Both the upstrap and `power.t.test()` tended to overestimate the power when compared to the true power estimates for small target sample sizes (20, 50; mean PE mostly between 1% and 7%) and showed great agreement for larger target sample sizes (≥ 80). In the case of power estimation for fixed effect size (first and second plots column), the lines spanning from the 25th to the 75th percentiles are very narrow for both approaches, indicating little variability of power estimates across the simulated samples.

**Figure 2:**
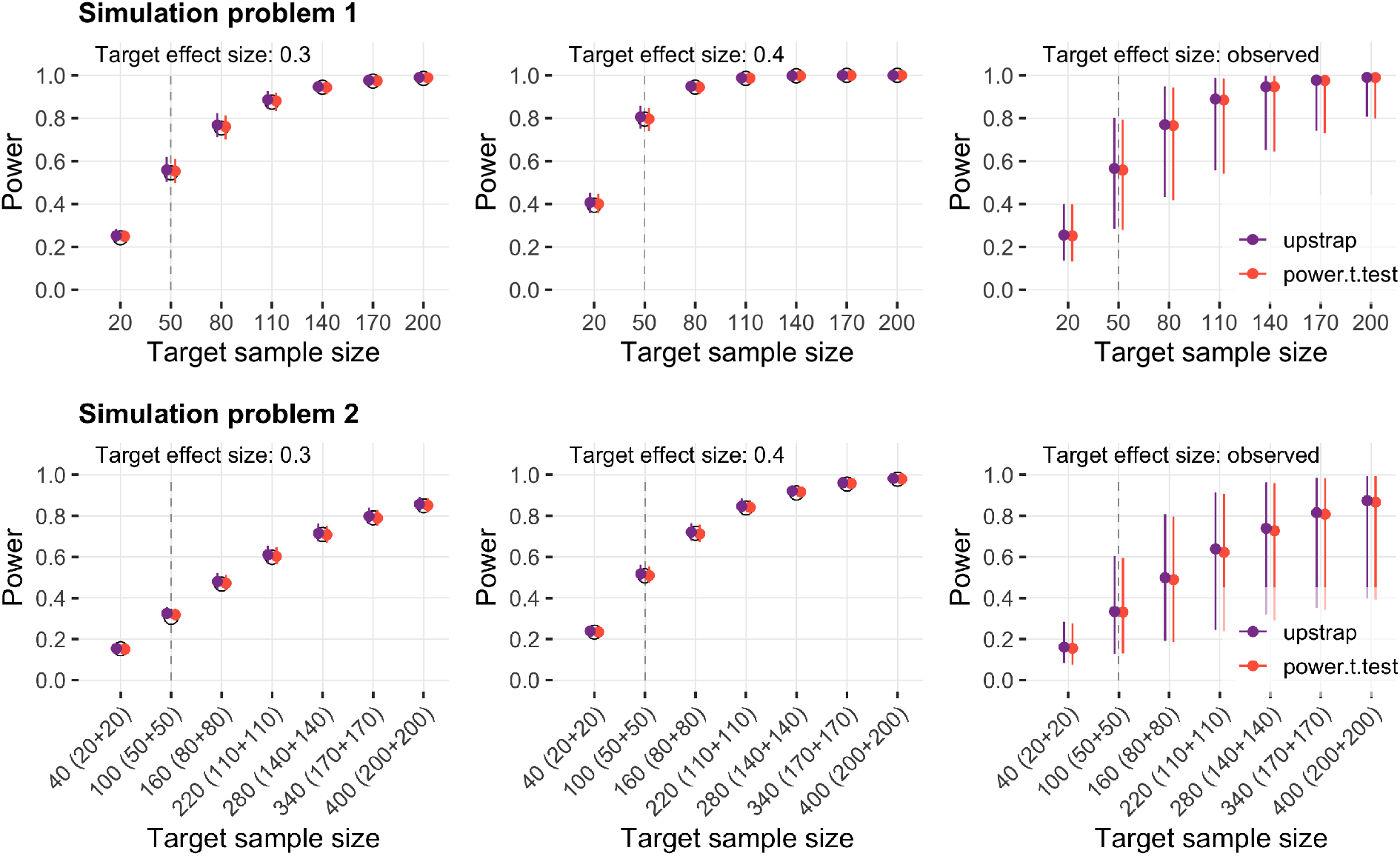
Power estimates (y-axis) at a range of target sample sizes (x-axis) in simulation problem 1 (one-sample t-test) and problem 2 (two-sample t-test). Colored points represent median and colored lines span from the 25th to the 75th percentiles of the power estimates from upstrap (purple color) and `power.t.test()` (orange color). Black circle points represent the true power estimate. Results are shown separately for three target effect size cases (plots columns 1,2 and 3, respectively). Vertical dashed grey lines denote the size of the observed sample.

#### 3.5.2 Simulation problems 3-6

Figure 3 displays the power estimates for simulation problems 3-6. The upstrap and `SIMR` approaches yielded very similar results across experiment repetitions, with absolute value of mean 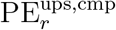 between them being below 2.6% across all experiment setups, and below 1% for most of them. In most cases, the upstrap/`SIMR` mean power estimates agree closely with the true power estimate; the largest discrepancies were present for simulation problem 4 (GLM with binary outcome) and problem 6 (GLMM with binary outcome) for the smallest target effect size considered (left plots column, 2nd and 4th plots rows).

**Figure 3:**
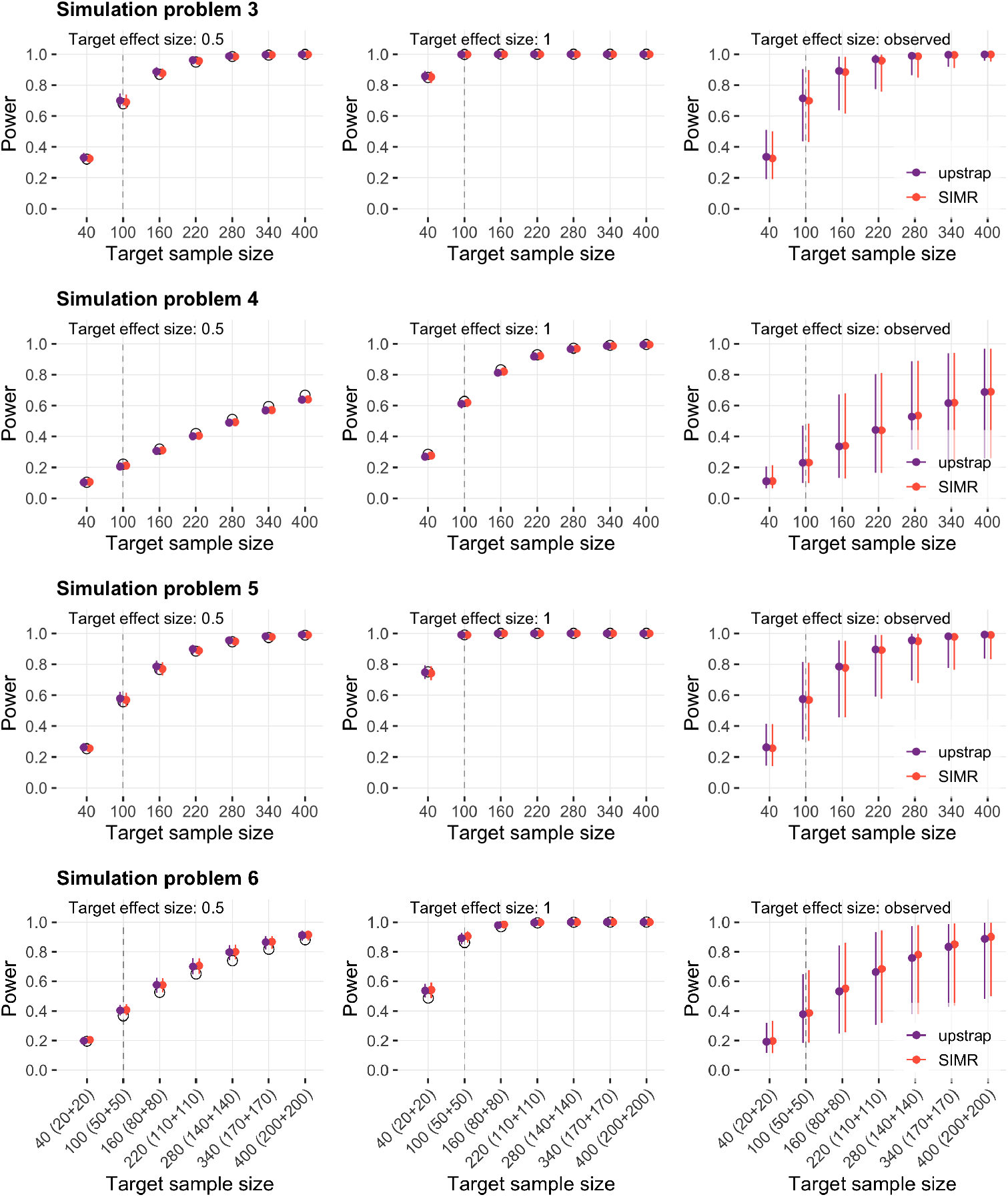
Power estimates (y-axis) at a range of target sample sizes (x-axis) in simulation problems 3-6: testing significance of coefficient in LM (problem 3), GLM with binary outcome (problem 4), LMM (problem 5), GLMM with binary outcome (problem 6), respectively. Values are showed for upstrap (purple color) and `SIMR` comparator (orange color) approaches. Black circle points represent the true power estimate. The lowest plots row shows split of target sample size across two levels of dichotomous covariate *X*_1_ (x-axis labels). Vertical dashed grey lines denote the size of the observed sample.

#### 3.5.3 Simulation problems 7-9

In simulation problems 7-9, upstrap was evaluated in a power estimation task in which the target distribution of a dichotomous covariate *X*_1_ (effect of interest) is changed compared to that from the observed data set. Table SM.2 in Supplementary Material shows a summary of power estimates and percentage errors (PEs) across simulation problems 7-9. Figure 4 shows power estimates at a range of target sample sizes in simulation problem 7 (two-sample t-test), problem 8 (testing significance of coefficient in linear model (LM)), and problem 9 (testing significance of coefficient in LMM). In each case, the proportion of *X*_1_ = 1 in an observed sample was 0.5, whereas the proportion of *X*_1_ = 1 in the target setup varied and equaled 0.3, 0.1 and 0.05 (plot columns 1, 2, and 3, respectively). Overall, upstrap demonstrated good to excellent alignment with true power estimates. The absolute value of mean PE between upstrap and true power was less than 4% for most of the cases.

**Figure 4:**
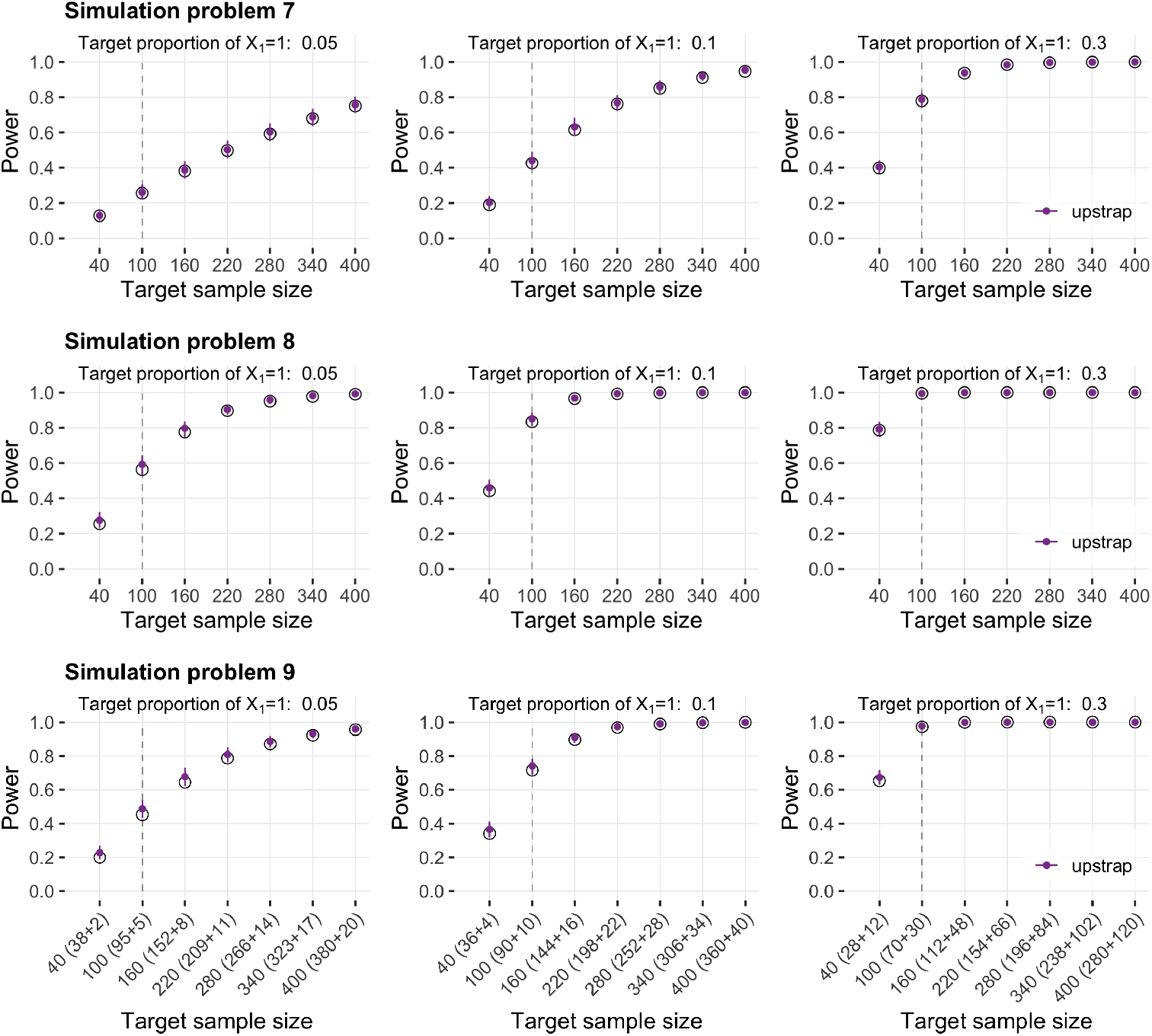
Power estimates (y-axis) at a range of target sample sizes (x-axis) in simulation problems 7-9: two-sample t-test (problem 7), testing significance of coefficient in LM (problem 8) and in LMM (problem 9), respectively. Values are showed for upstrap (purple color) and the true power estimate (black circle points). Results are showed across various values of target proportion of dichotomous covariate *X*_1_ equal 1 (0.05, 0.1 and 0.3 in plots column 1, 2, and 3, respectively). The lowest plots row shows split of the target sample size across two levels of dichotomous covariate *X*_1_ (x-axis labels). Vertical dashed grey lines denote the size of the observed sample.

#### 3.5.4 Simulation problems 10-12

In simulation problems 10-12, upstrap was evaluated in scenarios where the size of the observed sample was pushed to the lower limits. Table SM.3 in Supplementary Material shows a summary of power estimates and percentage errors (PEs) across simulation problems 10-12. Figure A.6 in Appendix A shows power estimates at a range of target sample sizes in simulation problem 10 (two-sample t-test), problem 11 (testing significance of coefficient in LM), and problem 12 (testing significance of coefficient in LMM). Four values of observed sample sizes were considered and equaled 10, 20, 40 and 100. Results are shown across different values of standard deviation assumed for the error term in data-generating models (plot columns 1, 2, and 3, respectively). Overall, upstrap demonstrated very good agreement with true power estimates for observed sample sizes 40 and 100, good agreement for observed sample size equal 20. Results show up to 40% of PE between upstrap and true power for the few most extreme cases considered in this problem (observed sample size equal 10 and target sample size on a lower end as well).

## 4 Application to a real data problem

The upstrap approach was used to estimate power of α-level tests in reanalysis of data from a cluster-randomized controlled trial on the impact of hotspot-targeted interventions on malaria transmission (2). Data are available online from the Dryad Digital Repository (https://bit.ly/3wrahch; (14)). R code of our reanalysis is available on GitHub (sub-directory URL: https://git.io/JsiP8).

### 4.1 Study description

(2) presented methods and findings from a cluster-randomized controlled trial conducted in western Kenya in 2012. In short, *N* =10 clusters of individuals were defined based on identification of 10 malaria hotspots, i.e. areas of serological and parasitological evidence of elevated levels of malaria transmission relative to their surrounding. Next, 5 clusters (consisting of total of 2, 082 individuals) were randomized into hotspot-targeted interventions arm and 5 clusters (2, 468 individuals) were randomized into placebo arm. The intervention consisted of a combination of “larviciding, distribution of long-lasting insecticide-treated nets, indoor residual spraying, and focal mass drug administration”. The placebo clusters followed Kenyan national policy of malaria control. The parasite prevalence was measured at three time points: baseline, 8 weeks, and 16 weeks post-intervention, across three sections defined based on the distance from the hotspot boundary: hotspot (0 meters), evaluation zone 1 (1–249 meters), evaluation zone 2 (250–500 meters).

The intervention effects were evaluated by comparing cluster-level parasite prevalence between control and intervention arms, separately for each of three evaluation zones. To make the comparison, first, an individual-level logistic regression model was fit with parasite status as an outcome and adjustment variables (baseline prevalence, age group, sex, altitude group, living in a house with open eaves) as covariates. Second, the model was used to estimate the expected outcome at the participant-level, and then to derive expected prevalence (average of participant-level outcomes) in each cluster. Third, cluster-level residuals (10 values) were calculated as the difference between the observed and expected prevalence. Fourth, the residuals of the control arm clusters and intervention arm clusters were compared using a t-test. Results from t-tests conducted on raw prevalence values without covariate adjustment were also reported. The paper concluded that the impact of interventions on parasite prevalence was “modest, transient, and restricted to the targeted hotspot areas”; it also stated that the trial “was not powered to detect subtle effects of hotspot-targeted interventions”. Data at participant-level resolution were deposited in the Dryad Digital Repository (14).

Below, we reanalyze the (2) data to replicate a subset of the main results. We then use the upstrap approach to estimate power trajectory of the relevant t-tests. Next, we perform an additional new analysis in which we use participant-level data in generalized estimating equations (GEE), and also perform an upstrap power trajectory estimation.

### 4.2 Replication of the study analysis results

A subset of the (2) main results regarding the intervention effect measured 8 weeks postintervention was considered. The assumed model was *Y_i_* = *β*_0_ + *β*_1_*X*_1*i*_ + *ϵ_i_*, where *i* = 1, …, *N*, *N* = 10 is the total number of clusters, *Y_i_* is the *i*-th cluster prevalence (unadjusted analysis) or *i*-th cluster residual of the observed minus expected prevalence (adjusted analysis), *X*_1*i*_ is the indicator whether the *i*-th cluster is in the intervention (*X*_1*i*_ = 1) or control (*X*_1*i*_ = 0) arm, 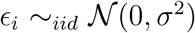. A two-sample t-test was used to test *H*_0_ : *β*_1_ = 0 against *H*_1_ : *β*_1_ ≠ 0, separately within each of three evaluation zones and for both unadjusted and adjusted analyses. In our replication analysis we relied on the R code provided by (15), who replicated the results from (2). Their R code is publicly available at https://osf.io/7ghfa/.

Table B.3 in Appendix B is an adaptation of “Table 3” from (2) comparing estimated parasite prevalence. In our reanalysis, the t-test rejected the null hypothesis of no treatment effect in hotspot area at the *α* = 0.05 level (p-value = 0.047). The hypothesis was not rejected for the other two evaluation zones (p-value equal 0.339 and 0.253, respectively).

### 4.3 Estimating power of a t-test using the upstrap

The upstrap approach was used to estimate power of three statistical tests corresponding to rows: 7, 14, and 21 in Table B.3 (adjusted analysis t-tests). A grid of target sample sizes {10, 12, 14, …, 60} was considered, and preserving the *X*_1*i*_ covariate distribution as in original sample (i.e., 50% cluster assignment to treatment and 50% clusters assignment to placebo arms) was assumed. *B* = 10, 000 upstrap replications were used. Top plots panel in Figure 5 displays estimated trajectories of t-test power. The number of clusters needed for achieving a power of 0.8 were estimated to be 18, 38, and 28 for the three area-specific tests, respectively. The results indicate that for the *N* =10 sample size available in the (2) paper’s study, the power of the test was 0.35, 0.14, and 0.19 for the three distance-based areas, respectively.

**Figure 5:**
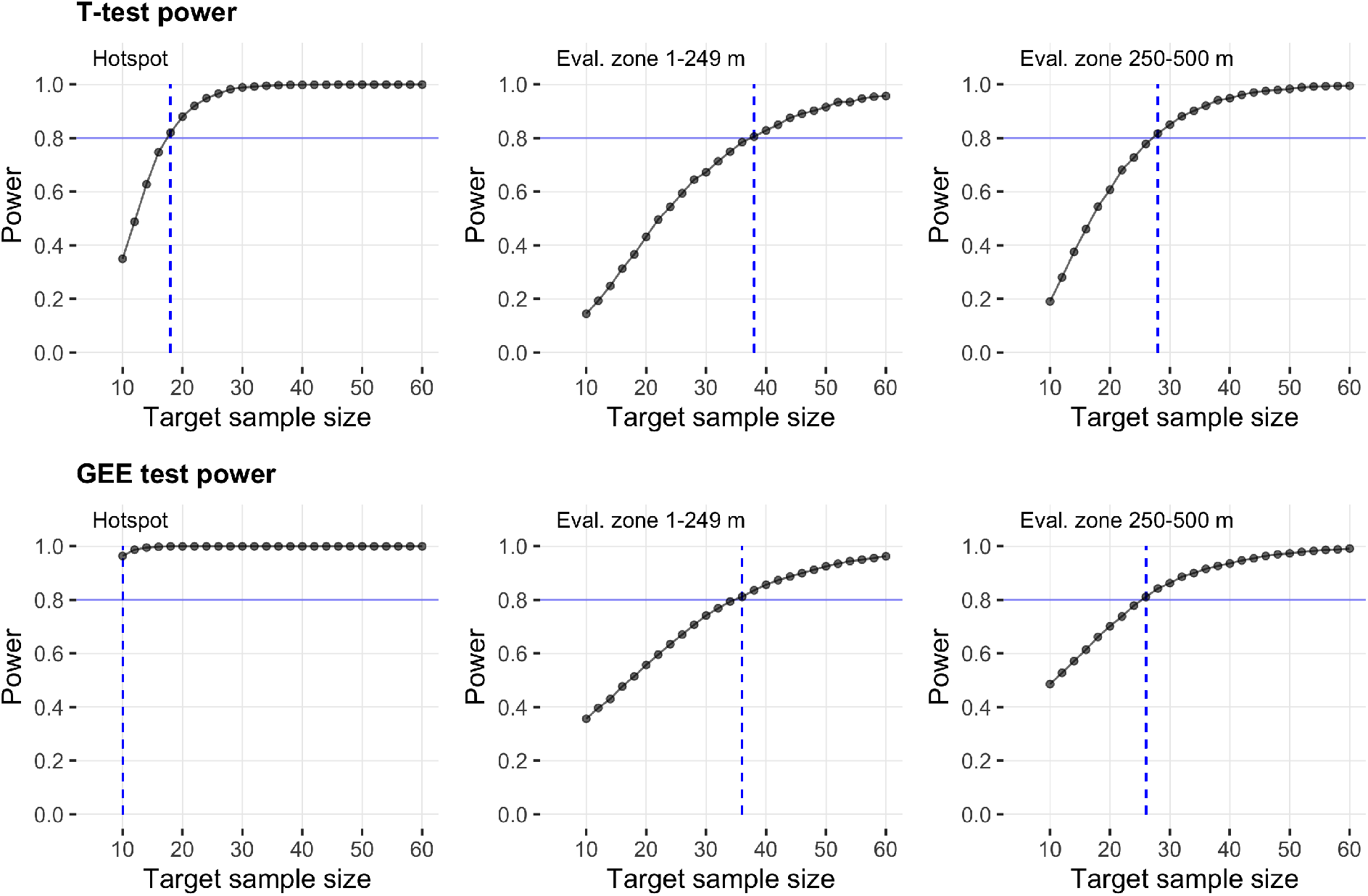
Top plots panel: estimated t-test power in the reanalysis of (2) paper’s data (adjusted t-test for a difference in prevalence between intervention and control groups at 8 weeks post-intervention). Bottom plots panel: estimated power in the additional analysis of (2) paper’s data (test of *β*_1_ coefficient from GEE model). The plots correspond to distance-based areas: hotspot, evaluation zone 1-249 meters, and evaluation zone 250-500 meters. Black dots show estimated power (y-axis) for a group sample size (x-axis). The horizontal solid blue line represents 0.8 power value; the vertical dashed blue line represents the smallest sample size for which the estimated power is at least 0.8.

**Figure 6:**
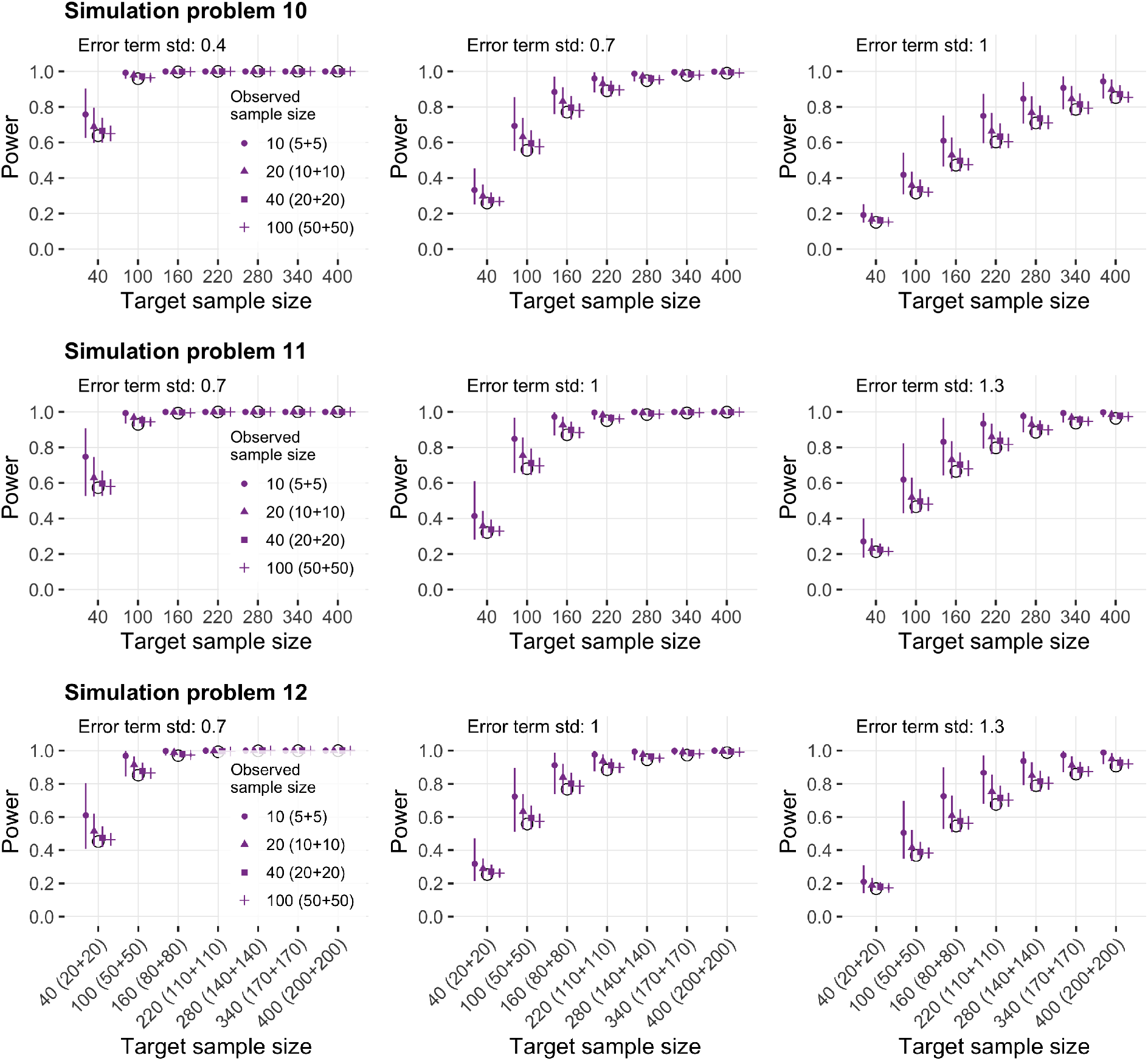
Power estimates (y-axis) at a range of target sample sizes (x-axis) in simulation problems 10-12: two-sample t-test (problem 10), testing significance of coefficient in LM (problem 11) and in LMM (problem 12), respectively. Values are showed for upstrap (purple color) and the true power estimate (black circle points). The four point shapes represent various observed sample size assumed in the experiment. Results are showed across various values of standard deviation assumed for error term in data-generating models.The lowest plots row shows split of the target sample size across two levels of dichotomous covariate *X*_1_ (x-axis labels).

### 4.4 Estimating power of a GEE-estimated effect test using the upstrap

Additional analysis of the (2) paper’s data was performed using participant-level data. The effect of the intervention was estimated via GEE, where the outcome is the recorded disease status (*Y_ij_* = 1 if positive and *Y_ij_* = 0 otherwise for *j*-th study participant in the *i*-th cluster). The linear predictor was modelled as a linear combination of intercept, cluster arm assignment *X*_1*i*_ (equal 1 if intervention and 0 if control) and participant’s predicted adjusted disease status *X*_*ij*2_ (equal 1 if positive and 0 otherwise; see Section 4.1 for details). The models were estimated using the `geeglm` function from `geepack` R package assuming a logit link function and exchangeable correlation structure within each cluster. In this model, the *β*_1_ coefficient of *X*_*i*1_ represents a population-averaged effect of the cluster-level intervention (when adjusted by the predicted status). We focused on the 8 weeks post-intervention effect was of interest, and three hotspot distance-based areas were considered separately.

Table B.4 in Appendix B summarizes the estimated 8 weeks post-intervention effect across three hotspot distance-based areas. In each case, the 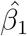 point estimate is negative; the statistical hypothesis *H*_0_ : *β*_1_ = 0 versus *H*_1_ : *β*_1_ ≠ 0 is rejected at significance level *α* = 0.05 for the hotspot (p-value < 0.0001), but not for either of the two evaluation zones (p-value equal 0.199 and 0.076, respectively).

The upstrap was used to estimate power of three statistical tests *H*_0_ : *β*_1_ = 0 versus *H*_1_ : *β*_1_ ≠ 0 from the new analysis. Similarly as earlier, a grid of target sample sizes {10, 12, 14, …, 60} was considered, and preserving the *X*_1*i*_ covariate distribution on subjectlevel as in the original sample was assumed. *B* = 10, 000 upstrap replications were used. Top plots panel in Figure 5 shows estimated trajectories of statistical power. The smallest group sample sizes for for which the estimated power is at least 0.8 were estimated as ≤ 10, 36, and 26 for the three area-specific tests, respectively: hotspot, evaluation zone 1-249 meters, evaluation zone 250-500 meters. For the second and third area-specific tests, the upstrap results using participant-level data therefore closely aligned with the ones for tests using cluster-level aggregate data in original analysis; for the test regarding the hotspot area, a smaller sample size was expected to be needed in our additional analysis (≤ 10 versus 18).

## 5 Discussion

Power estimation and sample size calculation are major components of statistical analyses. We proposed the upstrap-based methods to estimate power to detect both the effect size observed in the data and an effect size chosen by a researcher. The most important contribution of this paper is to evaluate the accuracy of the upstrap method for power estimation in simple and complex simulation scenarios.

First, our results indicate that for one- and two-sample t-test, the upstrap performs essentially identical to the well-established power estimation solutions. Second, in complex scenarios, the upstrap performed similarly the existing method from `SIMR` R package. Both approaches demonstrated very high agreement with the true power estimates. Notably, there is currently no feature to maintain balance or impose specific covariate class proportions in the `SIMR` R package. Such ability was demonstrated for upstrap in our simulations. It is unclear to us how such extension could be introduced for `SIMR` without employing upstrap multiple resampling idea. Third, the upstrap method is “read-and-use”, as it can be implemented by any analyst who is familiar with software allowing to (a) resample data, (b) run the statistical test of interest. To further facilitate the use by practitioners, in Supplementary Material, we demonstrate the upstrap method for estimating power in a series of examples. Fourth, the methodology is illustrated using a real dataset of a study with complex design. Fifth, the R code for our simulations and real data application example is publicaly available.

In addition, upstrap is particularly useful when the model might be misspecified, as re-sampling of independent units is done under weak modeling assumptions. For example, up-sampling and down-sampling independent units in a longitudinal data analysis will preserve the true correlation structures in the data. The `SIMR` approach requires a true model for the correlation structure.

Alas, our work is not without limitations. First, the upstrap computational time can be substantial. Steps that can speed up the process include employing parallel and/or cluster-based computing, as well as algorithmic improvements; for example, see our implementation of “rolling p-value” for one-sample t-test (https://git.io/JGqmh) and two-sample t-test (https://git.io/JGqOe). Second, this work has not covered a wide range of models, including longitudinal data or functional data regression models; further work is required to evaluate upstrap for power and sample size estimation in these settings.

In summary, the upstrap provides an easy to understand and implement solution for power estimation in complex models and scenarios. The examples given and the code provided will make the method easily available for practitioners.

## Appendices

### Appendix A Simulations

**Table A.2:**
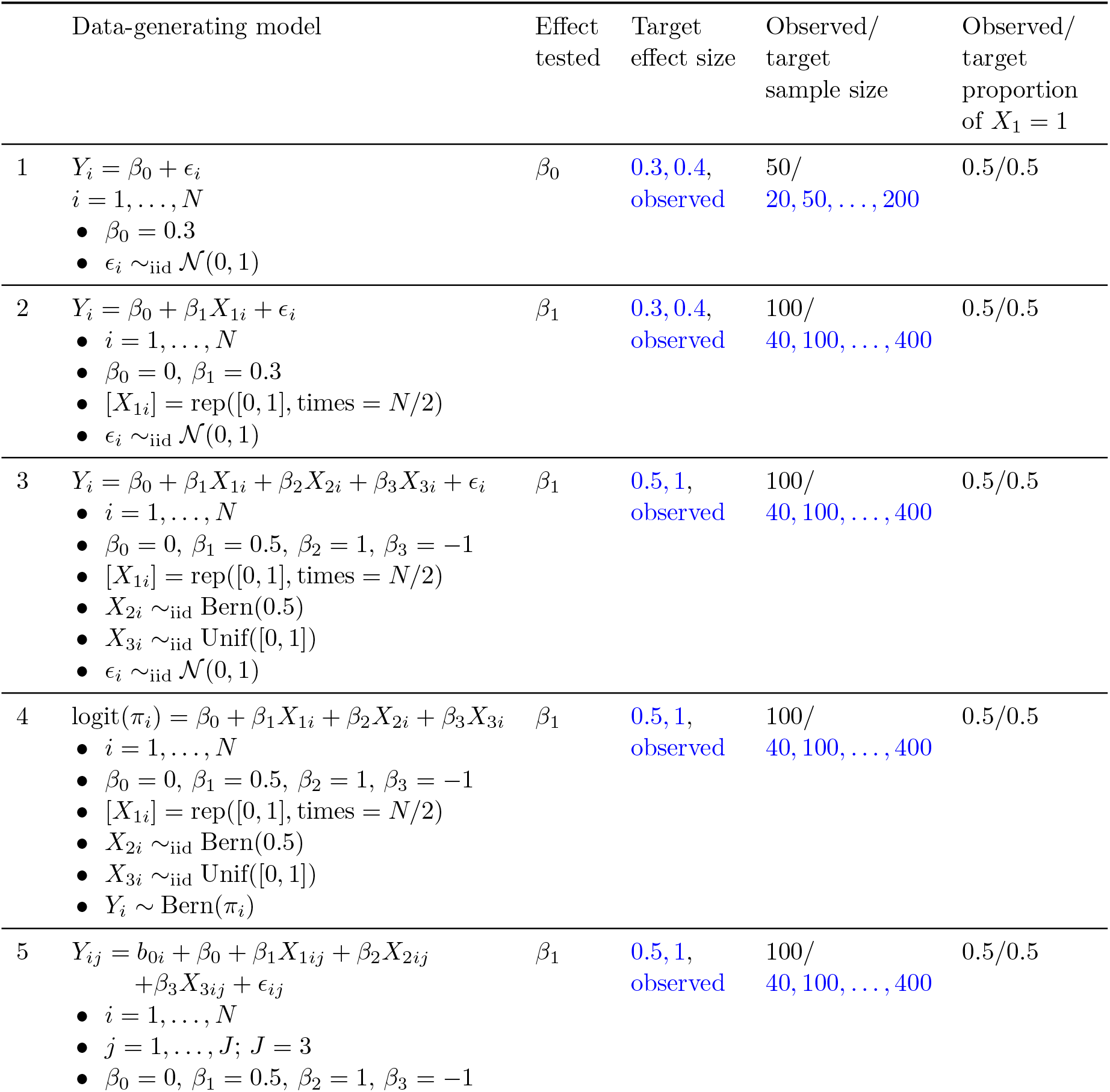
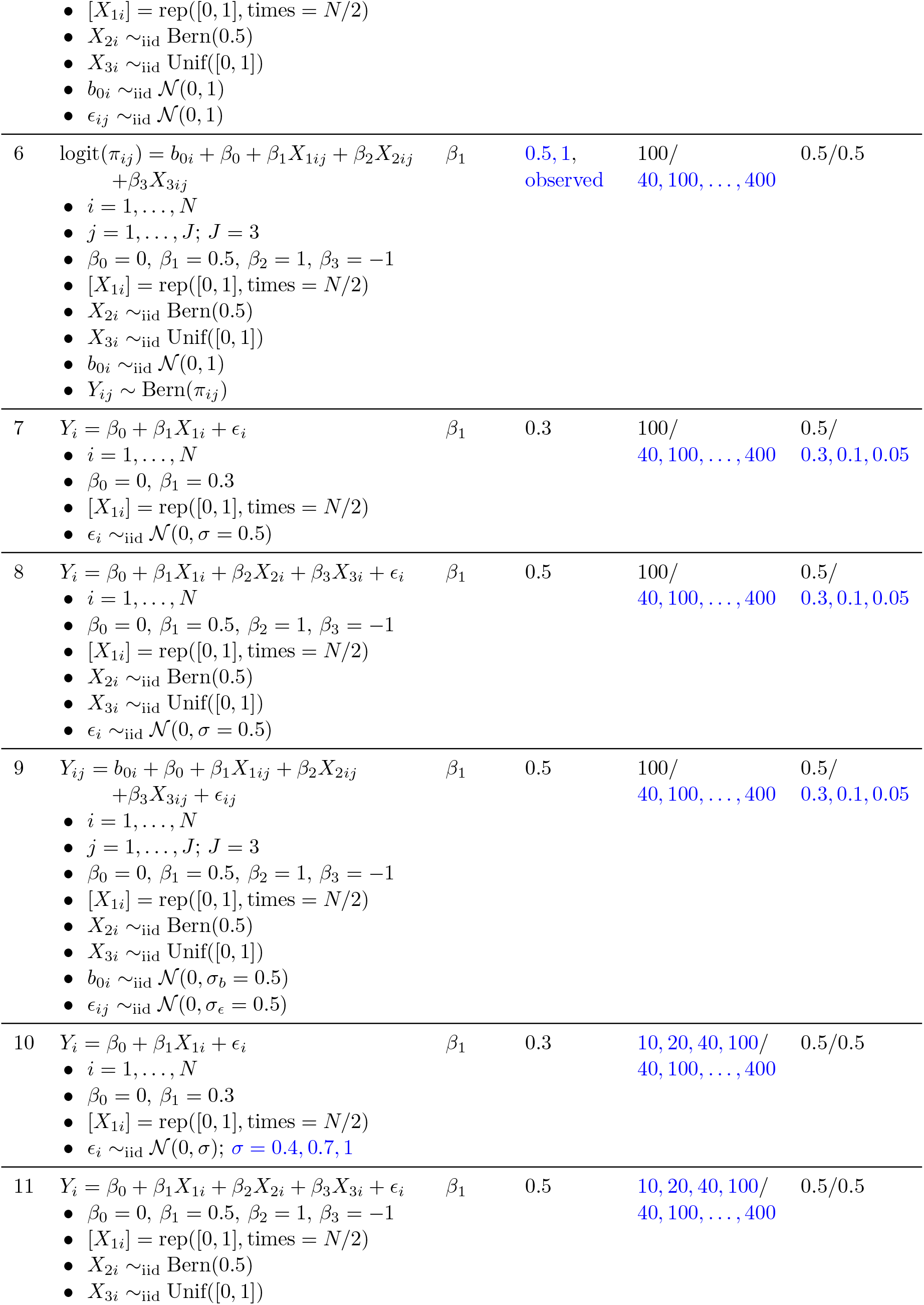
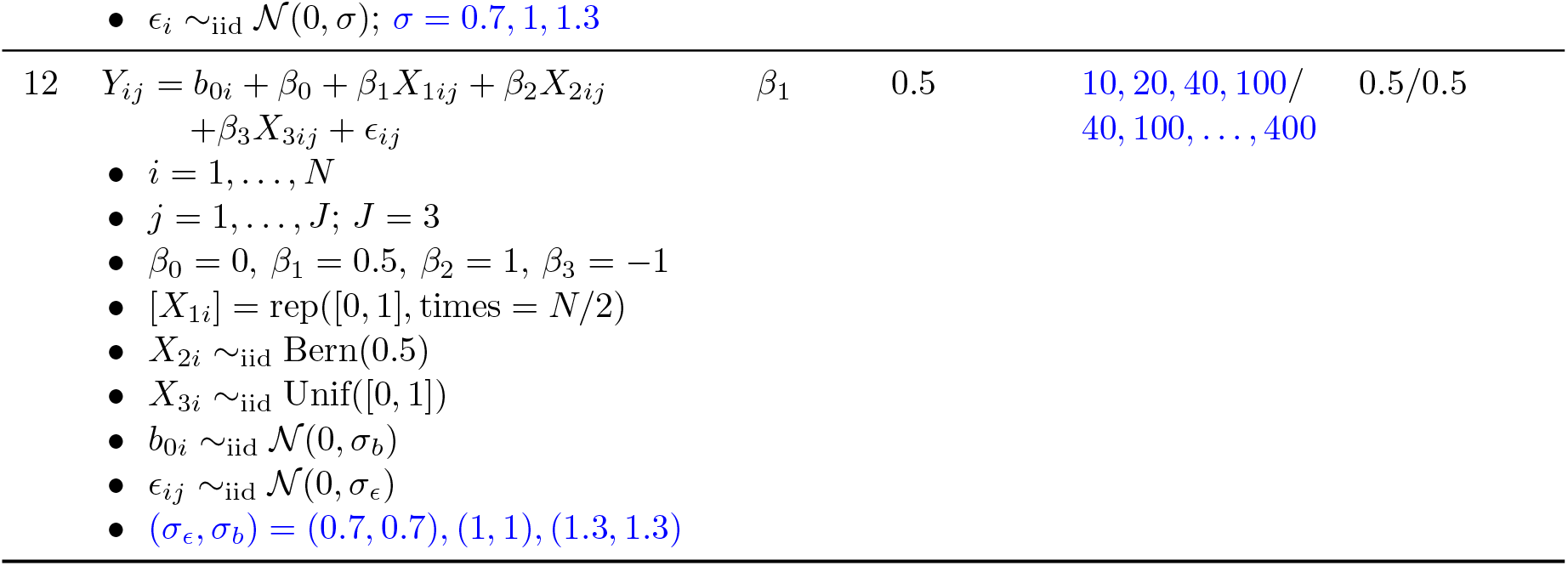
Summary of simulation setup across 12 different problems. In all cases, “sample size” refers to the number of independent units in the sample, indexed by *i*. In all cases, the covariates *X*_1*ij*_, *X*_2*ij*_ and *X*_3*ij*_ take the same value within a subject, i.e. *X_pij_* = *X_pi_* for *p* = 1, 2, 3 and for any *i* and *j*. For parameters whose values vary within a simulation problem, parameter’s values are displayed in blue color.

| | Data-generating model | Effect tested | Target effect size | Observed/<br>target<br>sample size | Observed/<br>target<br>proportion<br>of $X_1 = 1$ |
| --- | --- | --- | --- | --- | --- |
| 1 | $Y_i = \beta_0 + \epsilon_i$<br>$i = 1, \dots, N$<br>$\beta_0 = 0.3$<br>$\epsilon_i \sim_{\text{iid}} \mathcal{N}(0, 1)$ | $\beta_0$ | 0.3, 0.4,<br>observed | 50/<br>20, 50, ..., 200 | 0.5/0.5 |
| 2 | $Y_i = \beta_0 + \beta_1 X_{1i} + \epsilon_i$<br>$i = 1, \dots, N$<br>$\beta_0 = 0, \beta_1 = 0.3$<br>$[X_{1i}] = \text{rep}([0, 1], \text{times} = N/2)$<br>$\epsilon_i \sim_{\text{iid}} \mathcal{N}(0, 1)$ | $\beta_1$ | 0.3, 0.4,<br>observed | 100/<br>40, 100, ..., 400 | 0.5/0.5 |
| 3 | $Y_i = \beta_0 + \beta_1 X_{1i} + \beta_2 X_{2i} + \beta_3 X_{3i} + \epsilon_i$<br>$i = 1, \dots, N$<br>$\beta_0 = 0, \beta_1 = 0.5, \beta_2 = 1, \beta_3 = -1$<br>$[X_{1i}] = \text{rep}([0, 1], \text{times} = N/2)$<br>$X_{2i} \sim_{\text{iid}} \text{Bern}(0.5)$<br>$X_{3i} \sim_{\text{iid}} \text{Unif}([0, 1])$<br>$\epsilon_i \sim_{\text{iid}} \mathcal{N}(0, 1)$ | $\beta_1$ | 0.5, 1,<br>observed | 100/<br>40, 100, ..., 400 | 0.5/0.5 |
| 4 | $\text{logit}(\pi_i) = \beta_0 + \beta_1 X_{1i} + \beta_2 X_{2i} + \beta_3 X_{3i}$<br>$i = 1, \dots, N$<br>$\beta_0 = 0, \beta_1 = 0.5, \beta_2 = 1, \beta_3 = -1$<br>$[X_{1i}] = \text{rep}([0, 1], \text{times} = N/2)$<br>$X_{2i} \sim_{\text{iid}} \text{Bern}(0.5)$<br>$X_{3i} \sim_{\text{iid}} \text{Unif}([0, 1])$<br>$Y_i \sim \text{Bern}(\pi_i)$ | $\beta_1$ | 0.5, 1,<br>observed | 100/<br>40, 100, ..., 400 | 0.5/0.5 |
| 5 | $Y_{ij} = b_{0i} + \beta_0 + \beta_1 X_{1ij} + \beta_2 X_{2ij} + \beta_3 X_{3ij} + \epsilon_{ij}$<br>$i = 1, \dots, N$<br>$j = 1, \dots, J; J = 3$<br>$\beta_0 = 0, \beta_1 = 0.5, \beta_2 = 1, \beta_3 = -1$ | $\beta_1$ | 0.5, 1,<br>observed | 100/<br>40, 100, ..., 400 | 0.5/0.5 |
|  | • $[X_{1i}] = \text{rep}([0, 1], \text{times} = N/2)$<br>• $X_{2i} \sim_{\text{iid}} \text{Bern}(0.5)$<br>• $X_{3i} \sim_{\text{iid}} \text{Unif}([0, 1])$<br>• $b_{0i} \sim_{\text{iid}} \mathcal{N}(0, 1)$<br>• $\epsilon_{ij} \sim_{\text{iid}} \mathcal{N}(0, 1)$ |  |  |  |  |
| 6 | $\text{logit}(\pi_{ij}) = b_{0i} + \beta_0 + \beta_1 X_{1ij} + \beta_2 X_{2ij} + \beta_3 X_{3ij}$<br>• $i = 1, \dots, N$<br>• $j = 1, \dots, J; J = 3$<br>• $\beta_0 = 0, \beta_1 = 0.5, \beta_2 = 1, \beta_3 = -1$<br>• $[X_{1i}] = \text{rep}([0, 1], \text{times} = N/2)$<br>• $X_{2i} \sim_{\text{iid}} \text{Bern}(0.5)$<br>• $X_{3i} \sim_{\text{iid}} \text{Unif}([0, 1])$<br>• $b_{0i} \sim_{\text{iid}} \mathcal{N}(0, 1)$<br>• $Y_{ij} \sim \text{Bern}(\pi_{ij})$ | $\beta_1$ | 0.5, 1,<br>observed | 100/<br>40, 100, ..., 400 | 0.5/0.5 |
| 7 | $Y_i = \beta_0 + \beta_1 X_{1i} + \epsilon_i$<br>• $i = 1, \dots, N$<br>• $\beta_0 = 0, \beta_1 = 0.3$<br>• $[X_{1i}] = \text{rep}([0, 1], \text{times} = N/2)$<br>• $\epsilon_i \sim_{\text{iid}} \mathcal{N}(0, \sigma = 0.5)$ | $\beta_1$ | 0.3 | 100/<br>40, 100, ..., 400 | 0.5/<br>0.3, 0.1, 0.05 |
| 8 | $Y_i = \beta_0 + \beta_1 X_{1i} + \beta_2 X_{2i} + \beta_3 X_{3i} + \epsilon_i$<br>• $i = 1, \dots, N$<br>• $\beta_0 = 0, \beta_1 = 0.5, \beta_2 = 1, \beta_3 = -1$<br>• $[X_{1i}] = \text{rep}([0, 1], \text{times} = N/2)$<br>• $X_{2i} \sim_{\text{iid}} \text{Bern}(0.5)$<br>• $X_{3i} \sim_{\text{iid}} \text{Unif}([0, 1])$<br>• $\epsilon_i \sim_{\text{iid}} \mathcal{N}(0, \sigma = 0.5)$ | $\beta_1$ | 0.5 | 100/<br>40, 100, ..., 400 | 0.5/<br>0.3, 0.1, 0.05 |
| 9 | $Y_{ij} = b_{0i} + \beta_0 + \beta_1 X_{1ij} + \beta_2 X_{2ij} + \beta_3 X_{3ij} + \epsilon_{ij}$<br>• $i = 1, \dots, N$<br>• $j = 1, \dots, J; J = 3$<br>• $\beta_0 = 0, \beta_1 = 0.5, \beta_2 = 1, \beta_3 = -1$<br>• $[X_{1i}] = \text{rep}([0, 1], \text{times} = N/2)$<br>• $X_{2i} \sim_{\text{iid}} \text{Bern}(0.5)$<br>• $X_{3i} \sim_{\text{iid}} \text{Unif}([0, 1])$<br>• $b_{0i} \sim_{\text{iid}} \mathcal{N}(0, \sigma_b = 0.5)$<br>• $\epsilon_{ij} \sim_{\text{iid}} \mathcal{N}(0, \sigma_\epsilon = 0.5)$ | $\beta_1$ | 0.5 | 100/<br>40, 100, ..., 400 | 0.5/<br>0.3, 0.1, 0.05 |
| 10 | $Y_i = \beta_0 + \beta_1 X_{1i} + \epsilon_i$<br>• $i = 1, \dots, N$<br>• $\beta_0 = 0, \beta_1 = 0.3$<br>• $[X_{1i}] = \text{rep}([0, 1], \text{times} = N/2)$<br>• $\epsilon_i \sim_{\text{iid}} \mathcal{N}(0, \sigma); \sigma = 0.4, 0.7, 1$ | $\beta_1$ | 0.3 | 10, 20, 40, 100/<br>40, 100, ..., 400 | 0.5/0.5 |
| 11 | $Y_i = \beta_0 + \beta_1 X_{1i} + \beta_2 X_{2i} + \beta_3 X_{3i} + \epsilon_i$<br>• $\beta_0 = 0, \beta_1 = 0.5, \beta_2 = 1, \beta_3 = -1$<br>• $[X_{1i}] = \text{rep}([0, 1], \text{times} = N/2)$<br>• $X_{2i} \sim_{\text{iid}} \text{Bern}(0.5)$<br>• $X_{3i} \sim_{\text{iid}} \text{Unif}([0, 1])$ | $\beta_1$ | 0.5 | 10, 20, 40, 100/<br>40, 100, ..., 400 | 0.5/0.5 |
| | • $\epsilon_i \sim_{\text{iid}} \mathcal{N}(0, \sigma)$; $\sigma = 0.7, 1, 1.3$ | | | | |
| 12 | $Y_{ij} = b_{0i} + \beta_0 + \beta_1 X_{1ij} + \beta_2 X_{2ij}$<br>$\quad + \beta_3 X_{3ij} + \epsilon_{ij}$ | $\beta_1$ | 0.5 | 10, 20, 40, 100/<br>40, 100, …, 400 | 0.5/0.5 |
|  | • $i = 1, \dots, N$<br>• $j = 1, \dots, J$; $J = 3$<br>• $\beta_0 = 0, \beta_1 = 0.5, \beta_2 = 1, \beta_3 = -1$<br>• $[X_{1i}] = \text{rep}([0, 1], \text{times} = N/2)$<br>• $X_{2i} \sim_{\text{iid}} \text{Bern}(0.5)$<br>• $X_{3i} \sim_{\text{iid}} \text{Unif}([0, 1])$<br>• $b_{0i} \sim_{\text{iid}} \mathcal{N}(0, \sigma_b)$<br>• $\epsilon_{ij} \sim_{\text{iid}} \mathcal{N}(0, \sigma_\epsilon)$<br>• $(\sigma_\epsilon, \sigma_b) = (0.7, 0.7), (1, 1), (1.3, 1.3)$ |  |  |  |  |

### Appendix B Application to a real data problem

**Table B.3:** Adaptation of “Table 3” from (2) comparing estimated parasite prevalence in intervention and control clusters, across the two time points (baseline, 8 weeks postintervention), across three hotspot-specific areas (hotspot itself, evaluation zone 1-249 meters, evaluation zone 250-500 meters). Estimates and p-values are shown from: study’s original paper (2), previous work reanalysis (15), and this work reanalysis.

|  | Time point | Group or value | Bousema *and others* (2016) | Naudet *and others* (2018) | This paper’s reanalysis |
| --- | --- | --- | --- | --- | --- |
| nPCR Parasite Prevalence: |  |  |  |  |  |
| **Hotspot** |  |  |  |  |  |
| 1 | Baseline | Intervention | 20.1 |  | 21.4 |
| 2 | Baseline | Control | 18.6 |  | 18.6 |
| 3 | 8 weeks | Intervention | 9.2 |  | 9.2 |
| 4 | 8 weeks | Control | 19.4 |  | 19.4 |
| 5 | 8 weeks | Difference | 10.2 | 10.2 | 10.2 |
| 6 | 8 weeks | p-value | 0.075 | 0.095 | 0.095 |
| 7 | 8 weeks | Adj. p-value | 0.024 | 0.047 | 0.047 |
| **Eval. zone 1 (1-249 m)** |  |  |  |  |  |
| 8 | Baseline | Intervention | 13.2 |  | 13.2 |
| 9 | Baseline | Control | 11.7 |  | 11.7 |
| 10 | 8 weeks | Intervention | 12.3 |  | 12.3 |
| 11 | 8 weeks | Control | 15.9 |  | 15.9 |
| 12 | 8 weeks | Difference | 3.6 | 3.6 | 3.6 |
| 13 | 8 weeks | p-value | 0.219 | 0.2394 | 0.2394 |
| 14 | 8 weeks | Adj. p-value | 0.216 | 0.339 | 0.339 |
| **Eval. zone 2 (250-500 m)** |  |  |  |  |  |
| 15 | Baseline | Intervention | 13.5 |  | 13.5 |
| 16 | Baseline | Control | 16.9 |  | 16.9 |
| 17 | 8 weeks | Intervention | 9.1 |  | 9.1 |
| 18 | 8 weeks | Control | 12.9 |  | 13.1 |
| 19 | 8 weeks | Difference | 3.8 | 4 | 4 |
| 20 | 8 weeks | p-value | 0.199 | 0.1698 | 0.1698 |
| 21 | 8 weeks | Adj. p-value | 0.187 | 0.2533 | 0.2533 |

**Table B.4:**
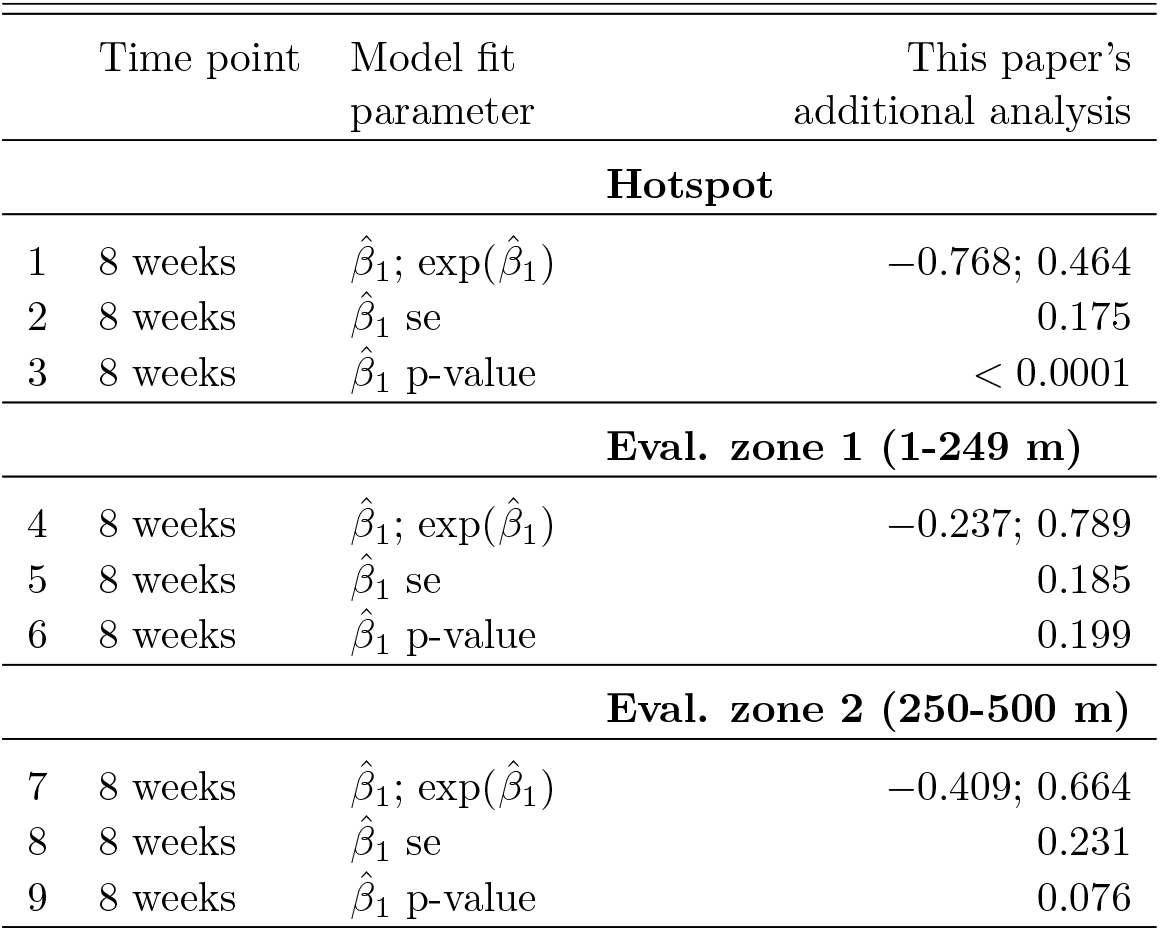
Estimated post-intervention effect across three hotspot distance-based areas using participant-level data. The *β*_1_ coefficient represents a population-averaged effect of the cluster-level intervention (when adjusted by the participant’s predicted status).

## Notes

https://datadryad.org/stash/dataset/doi:10.5061/dryad.nr8d8

### Competing Interest Statement

The authors have declared no competing interest.

### Summary of Updates

Added simulation experiments when the target covariate proportion is changed with respect to observed covariate proportion

