## SUPPLEMENTARY MATERIAL for "Upstrap for Estimating Power and Sample Size in Complex Models"

*To whom correspondence should be addressed.

### 1. Methods – R code examples

We demonstrate the upstrap method for estimating power in a series of examples accompanied by "read, adapt and use" R code. The problems include: one-sample t-test, two-sample t-test, testing significance of a coefficient in: linear regression, generalized linear regression with binary outcome, multilevel linear regression and generalized multilevel linear regression with binary outcome. All statistical tests assume significance level $\alpha = 0.05$.

#### 1.1 *One-sample t-test*

Consider a random sample from normal distribution. We are interested in estimating the power of the one-sample t-test.

We define simulation parameters and simulate a sample of size $N = 30$.

```
# simulation parameters
N <- 30
R_boot <- 1000
# simulate sample
set.seed(1)
x <- rnorm(n = N, mean = 0.3, sd = 1)
```

We get the observed effect size.

```
mean(x)
# [1] 0.3824582
```

We estimate test the power via upstrap.

- **Case**: target sample size $M = N = 30$, target effect size as observed in the sample ("observed power" case).

```
out <- rep(NA, R_boot)
for (rr in 1 : R_boot){
  x_rr <- sample(x, replace = TRUE)
  out[rr] <- (t.test(x_rr)$p.value < 0.05)
}
mean(out)
# [1] 0.593
```

- **Case**: target sample size $M = 40$, target effect size as observed in the sample.

```
out <- rep(NA, R_boot)
for (rr in 1 : R_boot){
  x_rr <- sample(x, size = 40, replace = TRUE)
  out[rr] <- (t.test(x_rr)$p.value < 0.05)
}
mean(out)
# [1] 0.716
```

- **Case**: target sample size $M = 40$, target effect size set to 0.5.

```
# update sample x to represent the target effect size
x_upd <- x + (0.5 - mean(x)) * 1
out <- rep(NA, R_boot)
for (rr in 1 : R_boot){
  x_rr <- sample(x_upd, size = 40, replace = TRUE)
  out[rr] <- (t.test(x_rr)$p.value < 0.05)
}
mean(out)
# [1] 0.917
```

##### 1.2 *Two-sample t-test*

Consider data for two groups (not paired) from normal distribution. We are interested in estimating power of two-sample t-test.

We define simulation parameters and simulate a sample of size (two groups combined) $N = 60$.

```
# simulation parameters
N <- 60
R_boot <- 1000
# simulate sample
set.seed(1)
x1 <- rnorm(N/2, mean = 0, sd = 1)
x2 <- rnorm(N/2, mean = 0.3, sd = 1)
```

We get the observed effect size.

```
mean(x2) - mean(x1)
# [1] 0.3503164
```

We estimate test power via upstrap.

- **Case**: target sample size $M = N = 60$, target effect size as observed in the sample ("observed power" case).

```
out <- rep(NA, R_boot)
for (rr in 1 : R_boot){
  x1_rr <- sample(x1, replace = TRUE)
  x2_rr <- sample(x2, replace = TRUE)
  pval_rr <- t.test(x1_rr, x2_rr, paired = FALSE, var.equal = TRUE)$p
     .value
  out[rr] <- (pval_rr < 0.05)
}
mean(out)
# [1] 0.349
```

- **Case**: target sample size $M = 80$, target effect size as observed in the sample.

```
M <- 80
out <- rep(NA, R_boot)
for (rr in 1 : R_boot){
  x1_rr <- sample(x1, size = M/2, replace = TRUE)
  x2_rr <- sample(x2, size = M/2, replace = TRUE)
  pval_rr <- t.test(x1_rr, x2_rr, paired = FALSE, var.equal = TRUE)$p
     .value
  out[rr] <- (pval_rr < 0.05)
}
mean(out)
# [1] 0.44
```

- **Case**: target sample size $M = 80$, target effect size set to 0.5.

```
M <- 80
# update sample to represent the target effect size
x1_upd <- x1
x2_upd <- x2 + (0.5 - (mean(x2) - mean(x1))) * 1
out <- rep(NA, R_boot)
for (rr in 1 : R_boot){
  x1_rr <- sample(x1_upd, size = M/2, replace = TRUE)
  x2_rr <- sample(x2_upd, size = M/2, replace = TRUE)
  pval_rr <- t.test(x1_rr, x2_rr, paired = FALSE, var.equal = TRUE)$p
     .value
  out[rr] <- (pval_rr < 0.05)
```

```
}
mean(out)
# [1] 0.752
```

##### 1.3 *Testing for significance of a coefficient in the linear model (LM)*

Consider a random sample with $N = 40$ independent observations (e.g., 40 subjects, 1 observation per subject). Assume a continuous outcome $Y$ and three covariates: dichotomous $X_1$, dichotomous $X_2$, continuous $X_3$. We are interested in estimating power of test for significance of the coefficient $\beta_1$ in linear model $Y_i = \beta_0 + \beta_1 X_{1i} + \beta_2 X_{2i} + \beta_3 X_{3i} + \varepsilon_i$, where $i = 1, \ldots, N$ and $\varepsilon_i \sim_{\text{iid}} N(0, \sigma^2)$.

We define simulation parameters and simulate a sample of size $N = 40$.

```
# simulation parameters
N       <- 40
coef_x0 <- 0
coef_x1 <- 0.6
coef_x2 <- 0.3
coef_x3 <- -0.1
sigma2  <- 1
R_boot  <- 1000
# simulate sample
set.seed(1)
subjid_i  <- 1 : N   # subject ID unique in data set
x1_i      <- rbinom(n = N, size = 1, prob = 0.5)
x2_i      <- rbinom(n = N, size = 1, prob = 0.5)
x3_i      <- runif(n = N, min = 18, max = 100)
eps_i     <- rnorm(N, sd = sqrt(sigma2))
y_i       <- coef_x0 + (coef_x1 * x1_i) + (coef_x2 * x2_i) + (coef_x3
     * x3_i) + eps_i
dat       <- data.frame(y = y_i, x1 = x1_i, x2 = x2_i, x3 = x3_i,
   subjid = subjid_i)
head(dat, 3)
# y           x1 x2       x3 subjid
# 1 -2.662590  0  1 53.64208      1
# 2 -7.381860  0  1 76.42620      2
# 3 -3.590214  1  1 50.79954      3
```

We get the observed effect size.

```
fit  <- lm(y ~ x1 + x2 + x3, data = dat)
coef(fit)["x1"]
# 0.3949275
```

We estimate the test power via the upstrap.

- **Case**: target sample size $M = N = 40$, target effect size as observed in the sample ("observed power" case).

```
out <- rep(NA, R_boot)
for (rr in 1 : R_boot){
  dat_rr_idx <- sample(1 : nrow(dat), replace = TRUE)
  dat_rr  <- dat[dat_rr_idx, ]
  fit_rr  <- lm(y ~ x1 + x2 + x3, data = dat_rr)
  pval_rr <- summary(fit_rr)$coef["x1", 4]
  out[rr] <- (pval_rr < 0.05)
}
mean(out)
# [1] 0.286
```

- **Case**: target sample size $M = 80$, target effect size as observed in the sample.

```
M <- 80
out <- rep(NA, R_boot)
for (rr in 1 : R_boot){
  dat_rr_idx <- sample(1 : nrow(dat), size = M, replace = TRUE)
  dat_rr  <- dat[dat_rr_idx, ]
  fit_rr  <- lm(y ~ x1 + x2 + x3, data = dat_rr)
  pval_rr <- summary(fit_rr)$coef["x1", 4]
  out[rr] <- (pval_rr < 0.05)
}
mean(out)
# [1] 0.511
```

- **Case**: target sample size $M = 80$, target effect size set to 0.5.

```
M <- 80
# update the outcome in the sample to represent the target effect
    size
dat_upd <- dat
dat_upd$y <- dat_upd$y + (0.5 - coef(fit)["x1"]) * dat_upd$x1
out <- rep(NA, R_boot)
for (rr in 1 : R_boot){
  dat_rr_idx <- sample(1 : nrow(dat_upd), size = M, replace = TRUE)
  dat_rr  <- dat_upd[dat_rr_idx, ]
  fit_rr  <- lm(y ~ x1 + x2 + x3, data = dat_rr)
  pval_rr <- summary(fit_rr)$coef["x1", 4]
```

```
  out[rr] <- (pval_rr < 0.05)
}
mean(out)
# [1] 0.686
```

##### 1.4 *Testing for significance of coefficient in generalized linear model (GLM)*

Consider a random sample with $N = 80$ independent observations (e.g. 80 subjects, 1 observation per subject). Assume binary outcome $Y$ and three covariates: dichotomous $X_1$, dichotomous $X_2$, continuous $X_3$. We are interested in estimating power of testing the coefficient $\beta_1$ in generalized linear model of a form $g(\pi_i) = \beta_0 + \beta_1 X_{1i} + \beta_2 X_{2i} + \beta_3 X_{3i}$, where $i = 1 \ldots, N$, $E(Y_i) = \pi_i$ and $g(\cdot)$ is a logit link function.

We define simulation parameters. We simulate a sample of size $N = 80$.

```
# simulation parameters
N       <- 80
coef_x0 <- -0.2
coef_x1 <- 0.5
coef_x2 <- 0.1
coef_x3 <- -0.01
sigma2  <- 1
R_boot  <- 1000
# simulate sample
set.seed(1)
subjid_i <- 1 : N
x1_i     <- rep(c(0, 1), times = N/2)
x2_i     <- rbinom(n = N, size = 1, prob = 0.5)
x3_i     <- runif(n = N, min = 18, max = 100)
XB_i     <- coef_x0 + (coef_x1 * x1_i) + (coef_x2 * x2_i) + (coef_x3
    * x3_i)
p_i      <- 1/(1 + exp(-XB_i))
y_i      <- rbinom(n = length(p_i), size = 1, prob = p_i)
dat      <- data.frame(y = y_i, x1 = x1_i, x2 = x2_i, x3 = x3_i,
    subjid = subjid_i)
head(dat, 3)
#   y x1 x2       x3 subjid
# 1 0  0  0 53.64208      1
# 2 0  1  0 76.42620      2
# 3 0  0  1 50.79954      3
```

We get the observed effect size.

```
fit  <- glm(y ~ x1 + x2 + x3, data = dat, family = binomial(link = "
    logit"))
coef(fit)["x1"]
# 0.7354476
```

We estimate test power via upstrap.

- **Case**: target sample size $M = N = 80$, target effect size as observed in the sample ("observed power" case).

```
out <- rep(NA, R_boot)
for (rr in 1 : R_boot){
  dat_rr_idx <- sample(1 : nrow(dat), replace = TRUE)
  dat_rr  <- dat[dat_rr_idx, ]
  fit_rr  <- glm(y ~ x1 + x2 + x3, data = dat_rr, family = binomial(
      link = "logit"))
  pval_rr <- summary(fit_rr)$coef[2, 4]
  out[rr] <- (pval_rr < 0.05)
}
mean(out)
# [1] 0.356
```

- **Case**: target sample size $M = 120$, target effect size as observed in the sample.

```
M <- 120
out <- rep(NA, R_boot)
for (rr in 1 : R_boot){
  dat_rr_idx <- sample(1 : nrow(dat), size = M, replace = TRUE)
  dat_rr  <- dat[dat_rr_idx, ]
  fit_rr  <- glm(y ~ x1 + x2 + x3, data = dat_rr, family = binomial(
      link = "logit"))
  pval_rr <- summary(fit_rr)$coef[2, 4]
  out[rr] <- (pval_rr < 0.05)
}
mean(out)
# [1] 0.441
```

- **Case**: target sample size $M = 120$, target effect size set to 1.0.

```
M <- 120
dat_upd <- dat
# define link term value, assuming target effect size
dat_upd$link_orig <- predict(fit, type = "link")
dat_upd$link_upd  <- dat_upd$link_orig  + (1 - coef(fit)["x1"]) * dat
   _upd$x1
dat_upd$res_upd   <- 1/(1 + exp(-dat_upd$link_upd))
out <- rep(NA, R_boot)
for (rr in 1 : R_boot){
  dat_rr_idx <- sample(1 : nrow(dat_upd), size = M, replace = TRUE)
  dat_rr    <- dat_upd[dat_rr_idx, ]
  # simulate response on resampled data, assuming target effect size
  dat_rr$y <- rbinom(n = nrow(dat_rr), size = 1, prob = dat_rr$res_
     upd)
  fit_rr    <- glm(y ~ x1 + x2 + x3, data = dat_rr, family = binomial(
     link = "logit"))
  pval_rr   <- summary(fit_rr)$coef[2, 4]
  out[rr]   <- (pval_rr < 0.05)
}
mean(out)
# [1] 0.757
```

##### 1.5 *Testing for significance of a coefficient in the linear mixed model (LMM)*

Consider a random sample with $N = 40$ subjects, $n_i = 3$ observations per each subject. Assume continuous outcome $Y$ and two covariates: dichotomous $X_1$, continuous $X_2$. We are interested in estimating power of test for significance of the subject-level (group-level) coefficient $\beta_1$ in linear mixed model $Y_{ij} = b_{0i} + \beta_0 + \beta_1 X_{1i} + \beta_2 X_{2i} + \epsilon_{ij}$, where $i = 1, \ldots, N$, $j = 1, 2, 3$, $b_{0i} \sim_{\text{iid}} \mathcal{N}(0, \tau^2)$, and $\varepsilon_i \sim_{\text{iid}} N(0, \sigma^2)$.

We define simulation parameters and simulate a sample of size $N = 40$.

```
# simulation parameters
N       <- 40
ni      <- 3
coef_x1 <- 1.2
coef_x2 <- 0.01
tau2    <- 1
sigma2  <- 1
R_boot  <- 1000
# simulate sample
```

```
set.seed(1)
subjid_i      <- 1:N                       # subject ID unique in
   data set
x1_i          <- rep(c(0, 1), times = N/2) # indicator of a "trial
   arm"
x2_i          <- runif(n = N, min = 18, max = 100)
b0_i          <- rnorm(n = N, mean = 0, sd = tau2)
eps_ij        <- rnorm(n = N * ni, mean = 0, sd = sigma2)
subjid_ij     <- rep(subjid_i, each = ni)
x1_ij         <- rep(x1_i, each = ni)
x2_ij         <- rep(x2_i, each = ni)
b0_ij         <- rep(b0_i, each = ni)
y_ij          <- b0_ij + coef_x1 * x1_ij + eps_ij
dat           <- data.frame(y = y_ij, x1 = x1_ij, x2 = x2_ij, subjid
   = subjid_ij)
head(dat, 4)
#           y x1       x2 subjid
# 1 3.3205951  0 39.77171      1
# 2 0.8797374  0 39.77171      1
# 3 1.6087167  0 39.77171      1
# 4 1.8101385  1 48.51416      2
```

We get the observed effect size.

```
library(lme4)
library(lmerTest)
fit  <- lmer(y ~ x1 + x2 + (1 | subjid), data = dat)
fixef(fit)["x1"]
# 0.6890758
```

We estimate test power via upstrap.

- **Case**: target sample size $M = N = 40$, target effect size as observed in the sample ("observed power" case).

```
out <- rep(NA, R_boot)
for (rr in 1 : R_boot){
  print(rr)
  dat_rr_subj_id <- sample(unique(dat$subjid), replace = TRUE)
  dat_rr  <- lapply(dat_rr_subj_id, function(subjid_tmp) dat[dat$
     subjid == subjid_tmp, ])
  dat_rr  <- do.call("rbind", dat_rr)
  # make new subject ID so as to treat subjects resampled >1 as
     unique ones
  dat_rr$subjid <- rep(1 : N, each = ni)
  fit_rr  <- lmer(y ~ x1 + x2 + (1 | subjid), data = dat_rr)
```

```
  pval_rr <- summary(fit_rr)$coef["x1", 5]
  out[rr] <- (pval_rr < 0.05)
}
mean(out)
# [1] 0.536
```

- **Case**: target sample size $M = 60$, target effect size as observed in the sample.

```
M <- 60
out <- rep(NA, R_boot)
for (rr in 1 : R_boot){
  print(rr)
  dat_rr_subj_id <- sample(unique(dat$subjid), size = M, replace =
     TRUE)
  dat_rr  <- lapply(dat_rr_subj_id, function(subjid_tmp) dat[dat$
     subjid == subjid_tmp, ])
  dat_rr  <- do.call("rbind", dat_rr)
  # make new subject ID so as to treat subjects resampled >1 as
     unique ones
  dat_rr$subjid <- rep(1 : M, each = ni)
  fit_rr  <- lmer(y ~ x1 + x2 + (1 | subjid), data = dat_rr)
  pval_rr <- summary(fit_rr)$coef["x1", 5]
  out[rr] <- (pval_rr < 0.05)
}
mean(out)
# [1] 0.714
```

- **Case**: target sample size $M = 60$, target effect size set to 1.2.

```
M <- 60
# target sample size: 60, target effect size: 1.2
dat_upd <- dat
dat_upd$y <- dat_upd$y + (1.2 - fixef(fit)["x1"]) * dat_upd$x1
out <- rep(NA, R_boot)
for (rr in 1 : R_boot){
  print(rr)
  dat_rr_subj_id <- sample(unique(dat_upd$subjid), size = M, replace
     = TRUE)
  dat_rr  <- lapply(dat_rr_subj_id, function(subjid_tmp) dat_upd[dat_
     upd$subjid == subjid_tmp, ])
  dat_rr  <- do.call("rbind", dat_rr)
  # make new subject ID so as to treat subjects resampled >1 as
     unique ones
  dat_rr$subjid <- rep(1 : M, each = ni)
```

```
  fit_rr  <- lmer(y ~ x1 + x2 + (1 | subjid), data = dat_rr)
  pval_rr <- summary(fit_rr)$coef["x1", 5]
  out[rr] <- (pval_rr < 0.05)
}
mean(out)
# [1] 0.994
```

##### 1.6 *Testing for significance of coefficient in generalized linear mixed model (GLMM)*

Consider a random sample with $N = 80$ subjects, $n_i = 3$ observations per each subject. Assume binary outcome $Y$ and two covariates: dichotomous $X_1$, continuous $X_2$. We are interested in estimating power of test for significance of the subject-level (group-level) coefficient $\beta_1$ in linear mixed model $g(\pi_{ij}) = b_{0i} + \beta_0 + \beta_1 X_{1i} + \beta_2 X_{2i}$, where $i = 1, \ldots, N$, $j = 1, 2, 3$, $b_{0i} \sim_{\text{iid}} \mathcal{N}(0, \tau^2)$, $E(Y_{ij}) = \pi_{ij}$ and $g(\cdot)$ is a logit link function.

We define simulation parameters. We simulate a sample of size $N = 80$.

```
# simulation parameters
N       <- 80
ni      <- 3
coef_x1 <- 0.8
coef_x2 <- 0.01
tau2    <- 1
sigma2  <- 1
R_boot  <- 1000
# simulate sample
set.seed(1)
subjid_i  <- 1:N # subject ID unique in data set
x1_i      <- rep(c(0, 1), times = N/2) # treatment arm
x2_i      <- runif(n = N, min = 18, max = 100)
b0_i      <- rnorm(n = N, mean = 0, sd = tau2)
subjid_ij <- rep(subjid_i, each = ni)
x1_ij     <- rep(x1_i, each = ni)
x2_ij     <- rep(x2_i, each = ni)
b0_ij     <- rep(b0_i, each = ni)
XB_ij     <- b0_ij + coef_x1 * x1_ij
p_ij      <- 1/(1 + exp(-XB_ij))
y_ij      <- rbinom(n = length(p_ij), size = 1, prob = p_ij)
dat       <- data.frame(y = y_ij, x1 = x1_ij, subjid = subjid_ij)
dat       <- data.frame(y = y_ij, x1 = x1_ij, x2 = x2_ij, subjid =
   subjid_ij)
head(dat, 4)
```

```
#    y x1        x2 subjid
# 1 0   0 39.77171      1
# 2 1   0 39.77171      1
# 3 1   0 39.77171      1
# 4 1   1 48.51416      2
```

We get the observed effect size.

```
library(lme4)
library(lmerTest)
fit  <- glmer(y ~ x1 + x2 + (1 | subjid), data = dat, family =
    binomial)
fixef(fit)["x1"]
# 0.8574488
```

We estimate test power via upstrap.

- **Case**: target sample size $M = N = 80$, target effect size as observed in the sample ("observed power" case).

```
out <- rep(NA, R_boot)
for (rr in 1 : R_boot){
  print(rr)
  dat_rr_subj_id <- sample(unique(dat$subjid), replace = TRUE)
  dat_rr  <- lapply(dat_rr_subj_id, function(subjid_tmp) dat[dat$
     subjid == subjid_tmp, ])
  dat_rr  <- do.call("rbind", dat_rr)
  # make new subject ID so as to treat subjects resampled >1 as
     unique ones
  dat_rr$subjid <- rep(1 : N, each = ni)
  fit_rr  <- glmer(y ~ x1 + x2 + (1 | subjid), data = dat_rr, family
     = binomial)
  pval_rr <- summary(fit_rr)$coef["x1", 4]
  out[rr] <- (pval_rr < 0.05)
}
mean(out)
# 0.652
```

- **Case**: target sample size $M = 100$, target effect size as observed in the sample.

```
M <- 100
out <- rep(NA, R_boot)
for (rr in 1 : R_boot){
  print(rr)
```

```
  dat_rr_subj_id <- sample(unique(dat$subjid), size = M, replace =
     TRUE)
  dat_rr  <- lapply(dat_rr_subj_id, function(subjid_tmp) dat[dat$
     subjid == subjid_tmp, ])
  dat_rr  <- do.call("rbind", dat_rr)
  # make new subject ID so as to treat subjects resampled >1 as
     unique ones
  dat_rr$subjid <- rep(1 : M, each = ni)
  fit_rr  <- glmer(y ~ x1 + x2 + (1 | subjid), data = dat_rr, family
     = binomial)
  pval_rr <- summary(fit_rr)$coef["x1", 4]
  out[rr] <- (pval_rr < 0.05)
}
mean(out)
# [1] 0.794
```

- **Case**: target sample size $M = 100$, target effect size set to 1.2.

```
M <- 100
set.seed(1)
out <- rep(NA, R_boot)
for (rr in 1 : R_boot){
  print(rr)
  dat_rr_subj_id <- sample(unique(dat$subjid), size = M, replace =
     TRUE)
  dat_rr  <- lapply(dat_rr_subj_id, function(subjid_tmp) dat[dat$
     subjid == subjid_tmp, ])
  dat_rr  <- do.call("rbind", dat_rr)
  # make new subject ID so as to treat subjects resampled >1 as
     unique ones
  dat_rr$subjid <- rep(1 : M, each = ni)
  # fit model, update coefficient for the size effect of interest
  fit_rr_sim_y  <- glmer(y ~ x1 + x2 + (1 | subjid), data = dat_rr,
     family = binomial)
  fixef(fit_rr_sim_y)["x1"] <- 1.2
  # simulate new outcome
  dat_rr$y <- simulate(fit_rr_sim_y, nsim = 1, newdata = dat_rr)[[1]]
  # fit "final" model
  fit_rr  <- glmer(y ~ x1 + x2 + (1 | subjid), data = dat_rr, family
     = binomial)
  pval_rr <- summary(fit_rr)$coef["x1", 4]
  out[rr] <- (pval_rr < 0.05)
}
mean(out)
# [1] 0.957
```

#### 2. Simulations

Table SM.1: Summary of power estimates and percentage errors (PEs) across simulation problems 1-6. The first and second table columns specify target effect size and target sample size. Table columns third to seventh show aggregates (mean and standard deviation) of power estimates and PE across $R = 1000$ independent experiment repetitions: $\hat{\text{pwr}}^{\text{ups}}$ – upstrap power, $\hat{\text{pwr}}^{\text{cmp}}$ – comparator approach power, $\text{PE}^{\text{ups,cmp}}$ – percentage error between upstrap and comparator, $\text{PE}^{\text{ups,true}}$ – percentage error between upstrap and true power, $\text{PE}^{\text{cmp,true}}$ – percentage error between comparator and true power.

**Simulation problem 1**

| | *Target effect size* | *Target sample size* | $\hat{\text{pwr}}^{\text{ups}}$ *mean (sd)* | $\hat{\text{pwr}}^{\text{cmp}}$ *mean (sd)* | $\text{PE}^{\text{ups,cmp}}$ *mean (sd)* | $\text{PE}^{\text{ups,true}}$ *mean (sd)* | $\text{PE}^{\text{cmp,true}}$ *mean (sd)* |
|---|---|---|---|---|---|---|---|
| 1 | 0.3 | 40 | 0.26 (0.05) | 0.25 (0.04) | 1.07 (9.36) | 5.95 (20.24) | 4.82 (17.62) |
| 2 | 0.3 | 100 | 0.57 (0.08) | 0.56 (0.08) | 1.33 (2.87) | 3.63 (15.23) | 2.29 (14.92) |
| 3 | 0.3 | 160 | 0.77 (0.08) | 0.76 (0.08) | 1.11 (1.96) | 1.71 (10.44) | 0.62 (10.36) |
| 4 | 0.3 | 220 | 0.88 (0.06) | 0.87 (0.06) | 0.73 (1.50) | 0.63 (6.70) | -0.08 (6.74) |
| 5 | 0.3 | 280 | 0.94 (0.04) | 0.94 (0.04) | 0.45 (1.05) | -0.40 (4.11) | -0.84 (4.18) |
| 6 | 0.3 | 340 | 0.97 (0.02) | 0.97 (0.02) | 0.26 (0.72) | -0.25 (2.47) | -0.50 (2.54) |
| 7 | 0.3 | 400 | 0.99 (0.01) | 0.98 (0.01) | 0.14 (0.47) | -0.02 (1.44) | -0.16 (1.50) |
| 8 | 0.4 | 40 | 0.41 (0.07) | 0.41 (0.07) | 0.72 (5.33) | 4.05 (18.15) | 3.28 (17.04) |
| 9 | 0.4 | 100 | 0.80 (0.07) | 0.79 (0.07) | 1.06 (2.03) | 0.80 (9.38) | -0.23 (9.31) |
| 10 | 0.4 | 160 | 0.94 (0.04) | 0.94 (0.04) | 0.45 (1.18) | -0.34 (4.12) | -0.78 (4.14) |
| 11 | 0.4 | 220 | 0.98 (0.02) | 0.98 (0.02) | 0.15 (0.56) | -0.27 (1.65) | -0.42 (1.67) |
| 12 | 0.4 | 280 | 1.00 (0.01) | 0.99 (0.01) | 0.06 (0.25) | -0.23 (0.62) | -0.29 (0.63) |
| 13 | 0.4 | 340 | 1.00 (0.00) | 1.00 (0.00) | 0.02 (0.14) | -0.09 (0.24) | -0.11 (0.23) |
| 14 | 0.4 | 400 | 1.00 (0.00) | 1.00 (0.00) | 0.00 (0.07) | -0.02 (0.10) | -0.02 (0.08) |
| 15 | observed | 40 | 0.29 (0.18) | 0.28 (0.18) | 5.23 (16.80) | - | - |
| 16 | observed | 100 | 0.54 (0.29) | 0.54 (0.29) | 3.09 (11.09) | - | - |
| 17 | observed | 160 | 0.67 (0.31) | 0.67 (0.31) | 2.06 (8.78) | - | - |
| 18 | observed | 220 | 0.74 (0.30) | 0.74 (0.30) | 1.76 (7.82) | - | - |
| 19 | observed | 280 | 0.79 (0.29) | 0.79 (0.29) | 1.34 (7.18) | - | - |
| 20 | observed | 340 | 0.82 (0.28) | 0.82 (0.28) | 1.40 (7.91) | - | - |
| 21 | observed | 400 | 0.84 (0.27) | 0.84 (0.27) | 1.30 (7.84) | - | - |

**Simulation problem 2**

| | *Target effect size* | *Target sample size* | $\hat{\text{pwr}}^{\text{ups}}$ *mean (sd)* | $\hat{\text{pwr}}^{\text{cmp}}$ *mean (sd)* | $\text{PE}^{\text{ups,cmp}}$ *mean (sd)* | $\text{PE}^{\text{ups,true}}$ *mean (sd)* | $\text{PE}^{\text{cmp,true}}$ *mean (sd)* |
|---|---|---|---|---|---|---|---|
| 1 | 0.3 | 40 | 0.16 (0.02) | 0.15 (0.02) | 2.73 (9.35) | 2.09 (13.86) | -0.57 (10.38) |
| 2 | 0.3 | 100 | 0.33 (0.04) | 0.32 (0.04) | 1.66 (4.88) | 6.75 (13.62) | 5.03 (12.60) |
| 3 | 0.3 | 160 | 0.48 (0.06) | 0.48 (0.05) | 1.61 (3.25) | 3.26 (12.08) | 1.64 (11.55) |
| 4 | 0.3 | 220 | 0.61 (0.06) | 0.60 (0.06) | 1.45 (2.44) | 2.55 (10.38) | 1.10 (10.17) |
| 5 | 0.3 | 280 | 0.72 (0.06) | 0.71 (0.06) | 1.24 (1.99) | 0.99 (8.72) | -0.23 (8.57) |
| 6 | 0.3 | 340 | 0.80 (0.06) | 0.79 (0.06) | 1.08 (1.67) | 0.75 (7.15) | -0.32 (7.12) |
| 7 | 0.3 | 400 | 0.86 (0.05) | 0.85 (0.05) | 0.88 (1.37) | 0.70 (5.78) | -0.16 (5.81) |
| 8 | 0.4 | 40 | 0.24 (0.03) | 0.24 (0.03) | 1.52 (7.14) | 2.92 (14.05) | 1.37 (11.85) |

| | | | | | | | |
|---|---|---|---|---|---|---|---|
| 9 | 0.4 | 100 | 0.52 (0.06) | 0.51 (0.06) | 1.48 (3.11) | 2.50 (11.64) | 1.01 (11.16) |
| 10 | 0.4 | 160 | 0.72 (0.06) | 0.71 (0.06) | 1.14 (2.04) | 0.78 (8.62) | -0.34 (8.49) |
| 11 | 0.4 | 220 | 0.85 (0.05) | 0.84 (0.05) | 0.88 (1.42) | 0.93 (6.07) | 0.06 (6.04) |
| 12 | 0.4 | 280 | 0.92 (0.04) | 0.91 (0.04) | 0.52 (0.99) | 0.67 (4.04) | 0.15 (4.05) |
| 13 | 0.4 | 340 | 0.96 (0.02) | 0.95 (0.02) | 0.35 (0.71) | 0.29 (2.55) | -0.06 (2.60) |
| 14 | 0.4 | 400 | 0.98 (0.02) | 0.98 (0.02) | 0.23 (0.50) | -0.02 (1.60) | -0.25 (1.61) |
| 15 | observed | 40 | 0.21 (0.15) | 0.20 (0.15) | 11.71 (23.27) | - | - |
| 16 | observed | 100 | 0.39 (0.28) | 0.38 (0.28) | 7.61 (19.74) | - | - |
| 17 | observed | 160 | 0.50 (0.32) | 0.50 (0.32) | 6.16 (17.40) | - | - |
| 18 | observed | 220 | 0.58 (0.34) | 0.57 (0.34) | 5.38 (16.53) | - | - |
| 19 | observed | 280 | 0.63 (0.34) | 0.62 (0.35) | 4.80 (15.17) | - | - |
| 20 | observed | 340 | 0.67 (0.34) | 0.66 (0.35) | 3.88 (13.42) | - | - |
| 21 | observed | 400 | 0.70 (0.34) | 0.69 (0.34) | 3.57 (12.89) | - | - |

**Simulation problem 3**

| | *Target effect size* | *Target sample size* | $\hat{\text{pwr}}^{\text{ups}}$ *mean (sd)* | $\hat{\text{pwr}}^{\text{cmp}}$ *mean (sd)* | $\text{PE}^{\text{ups,cmp}}$ *mean (sd)* | $\text{PE}^{\text{ups,true}}$ *mean (sd)* | $\text{PE}^{\text{cmp,true}}$ *mean (sd)* |
|---|---|---|---|---|---|---|---|
| 1 | 0.5 | 40 | 0.33 (0.04) | 0.33 (0.05) | 1.70 (8.61) | 3.38 (13.90) | 2.09 (14.32) |
| 2 | 0.5 | 100 | 0.70 (0.06) | 0.69 (0.07) | 1.06 (3.20) | 2.81 (9.55) | 1.80 (9.61) |
| 3 | 0.5 | 160 | 0.88 (0.05) | 0.87 (0.05) | 1.10 (1.87) | 1.36 (5.26) | 0.29 (5.48) |
| 4 | 0.5 | 220 | 0.96 (0.03) | 0.95 (0.03) | 0.57 (1.07) | 0.74 (2.66) | 0.18 (2.77) |
| 5 | 0.5 | 280 | 0.99 (0.01) | 0.98 (0.01) | 0.26 (0.60) | 0.01 (1.22) | -0.25 (1.34) |
| 6 | 0.5 | 340 | 0.99 (0.01) | 0.99 (0.01) | 0.11 (0.37) | 0.06 (0.55) | -0.06 (0.62) |
| 7 | 0.5 | 400 | 1.00 (0.00) | 1.00 (0.00) | 0.05 (0.23) | -0.04 (0.25) | -0.09 (0.30) |
| 8 | 1 | 40 | 0.85 (0.05) | 0.85 (0.06) | 0.77 (3.48) | 0.52 (5.96) | -0.15 (6.52) |
| 9 | 1 | 100 | 1.00 (0.00) | 1.00 (0.00) | 0.02 (0.24) | -0.06 (0.30) | -0.08 (0.32) |
| 10 | 1 | 160 | 1.00 (0.00) | 1.00 (0.00) | -0.00 (0.03) | -0.00 (0.02) | -0.00 (0.02) |
| 11 | 1 | 220 | 1.00 (0.00) | 1.00 (0.00) | 0.00 (0.01) | 0.00 (0.00) | -0.00 (0.01) |
| 12 | 1 | 280 | 1.00 (0.00) | 1.00 (0.00) | 0.00 (0.00) | 0.00 (0.00) | 0.00 (0.00) |
| 13 | 1 | 340 | 1.00 (0.00) | 1.00 (0.00) | 0.00 (0.00) | 0.00 (0.00) | 0.00 (0.00) |
| 14 | 1 | 400 | 1.00 (0.00) | 1.00 (0.00) | 0.00 (0.00) | 0.00 (0.00) | 0.00 (0.00) |
| 15 | observed | 40 | 0.36 (0.21) | 0.36 (0.21) | 2.19 (10.79) | - | - |
| 16 | observed | 100 | 0.65 (0.28) | 0.64 (0.28) | 1.15 (6.76) | - | - |
| 17 | observed | 160 | 0.77 (0.26) | 0.77 (0.26) | 1.35 (5.53) | - | - |
| 18 | observed | 220 | 0.84 (0.24) | 0.83 (0.24) | 1.02 (4.41) | - | - |
| 19 | observed | 280 | 0.87 (0.22) | 0.87 (0.22) | 0.87 (4.11) | - | - |
| 20 | observed | 340 | 0.90 (0.20) | 0.89 (0.21) | 0.69 (3.74) | - | - |
| 21 | observed | 400 | 0.92 (0.19) | 0.91 (0.19) | 0.71 (4.18) | - | - |

**Simulation problem 4**

| | *Target effect size* | *Target sample size* | $\hat{\text{pwr}}^{\text{ups}}$ *mean (sd)* | $\hat{\text{pwr}}^{\text{cmp}}$ *mean (sd)* | $\text{PE}^{\text{ups,cmp}}$ *mean (sd)* | $\text{PE}^{\text{ups,true}}$ *mean (sd)* | $\text{PE}^{\text{cmp,true}}$ *mean (sd)* |
|---|---|---|---|---|---|---|---|
| 1 | 0.5 | 40 | 0.10 (0.01) | 0.10 (0.02) | -0.61 (17.43) | -1.32 (13.11) | 1.06 (15.37) |
| 2 | 0.5 | 100 | 0.21 (0.02) | 0.21 (0.02) | -1.91 (8.25) | -6.38 (8.29) | -4.23 (8.37) |
| 3 | 0.5 | 160 | 0.31 (0.02) | 0.31 (0.02) | -0.83 (6.79) | -4.06 (7.67) | -3.06 (7.45) |
| 4 | 0.5 | 220 | 0.40 (0.03) | 0.40 (0.03) | -0.54 (5.36) | -4.48 (6.89) | -3.83 (6.82) |

| | | | | | | | |
|---|---|---|---|---|---|---|---|
| 5 | 0.5 | 280 | 0.49 (0.03) | 0.49 (0.03) | -0.39 (4.51) | -4.69 (6.38) | -4.23 (6.22) |
| 6 | 0.5 | 340 | 0.57 (0.03) | 0.57 (0.03) | -0.40 (3.90) | -4.87 (5.78) | -4.42 (5.74) |
| 7 | 0.5 | 400 | 0.63 (0.04) | 0.64 (0.03) | -0.47 (3.46) | -4.84 (5.38) | -4.35 (5.20) |
| 8 | 1 | 40 | 0.27 (0.03) | 0.27 (0.04) | -1.79 (11.73) | -6.66 (11.64) | -4.08 (13.62) |
| 9 | 1 | 100 | 0.61 (0.04) | 0.62 (0.04) | -1.36 (3.62) | -3.12 (6.41) | -1.71 (6.54) |
| 10 | 1 | 160 | 0.81 (0.04) | 0.81 (0.04) | -0.60 (2.23) | -2.62 (4.23) | -2.02 (4.23) |
| 11 | 1 | 220 | 0.91 (0.02) | 0.92 (0.02) | -0.31 (1.35) | -1.56 (2.58) | -1.24 (2.60) |
| 12 | 1 | 280 | 0.96 (0.01) | 0.96 (0.01) | -0.13 (0.87) | -0.87 (1.52) | -0.74 (1.50) |
| 13 | 1 | 340 | 0.98 (0.01) | 0.99 (0.01) | -0.06 (0.58) | -0.56 (0.83) | -0.50 (0.83) |
| 14 | 1 | 400 | 0.99 (0.00) | 0.99 (0.00) | -0.03 (0.35) | -0.17 (0.46) | -0.14 (0.44) |
| 15 | observed | 40 | 0.15 (0.12) | 0.16 (0.12) | -0.22 (16.72) | - | - |
| 16 | observed | 100 | 0.31 (0.25) | 0.32 (0.25) | -0.29 (11.79) | - | - |
| 17 | observed | 160 | 0.41 (0.30) | 0.41 (0.30) | 0.10 (10.26) | - | - |
| 18 | observed | 220 | 0.48 (0.33) | 0.49 (0.33) | 0.05 (9.13) | - | - |
| 19 | observed | 280 | 0.54 (0.34) | 0.54 (0.34) | -0.12 (8.98) | - | - |
| 20 | observed | 340 | 0.58 (0.35) | 0.58 (0.35) | 0.09 (7.93) | - | - |
| 21 | observed | 400 | 0.61 (0.35) | 0.61 (0.35) | 0.27 (7.05) | - | - |

**Simulation problem 5**

| | *Target effect size* | *Target sample size* | $\hat{\text{pwr}}^{\text{ups}}$ *mean (sd)* | $\hat{\text{pwr}}^{\text{cmp}}$ *mean (sd)* | $\text{PE}^{\text{ups,cmp}}$ *mean (sd)* | $\text{PE}^{\text{ups,true}}$ *mean (sd)* | $\text{PE}^{\text{cmp,true}}$ *mean (sd)* |
|---|---|---|---|---|---|---|---|
| 1 | 0.5 | 40 | 0.26 (0.04) | 0.26 (0.04) | 2.37 (10.20) | 4.13 (14.85) | 2.18 (14.42) |
| 2 | 0.5 | 100 | 0.58 (0.06) | 0.57 (0.06) | 1.51 (3.92) | 3.71 (11.60) | 2.26 (11.61) |
| 3 | 0.5 | 160 | 0.78 (0.06) | 0.77 (0.06) | 1.67 (2.56) | 1.90 (7.94) | 0.28 (8.06) |
| 4 | 0.5 | 220 | 0.89 (0.04) | 0.88 (0.05) | 1.11 (1.72) | 1.04 (5.04) | -0.05 (5.16) |
| 5 | 0.5 | 280 | 0.95 (0.03) | 0.94 (0.03) | 0.72 (1.13) | 0.52 (2.97) | -0.18 (3.15) |
| 6 | 0.5 | 340 | 0.98 (0.02) | 0.97 (0.02) | 0.42 (0.77) | 0.42 (1.69) | 0.01 (1.87) |
| 7 | 0.5 | 400 | 0.99 (0.01) | 0.99 (0.01) | 0.24 (0.52) | 0.13 (0.94) | -0.11 (1.06) |
| 8 | 1 | 40 | 0.75 (0.06) | 0.74 (0.07) | 1.11 (4.37) | -0.52 (8.48) | -1.48 (8.89) |
| 9 | 1 | 100 | 0.99 (0.01) | 0.99 (0.01) | 0.10 (0.58) | -0.28 (1.10) | -0.39 (1.14) |
| 10 | 1 | 160 | 1.00 (0.00) | 1.00 (0.00) | 0.01 (0.11) | -0.01 (0.11) | -0.02 (0.12) |
| 11 | 1 | 220 | 1.00 (0.00) | 1.00 (0.00) | 0.00 (0.02) | -0.00 (0.02) | -0.00 (0.02) |
| 12 | 1 | 280 | 1.00 (0.00) | 1.00 (0.00) | -0.00 (0.01) | -0.00 (0.01) | -0.00 (0.00) |
| 13 | 1 | 340 | 1.00 (0.00) | 1.00 (0.00) | 0.00 (0.00) | 0.00 (0.00) | 0.00 (0.00) |
| 14 | 1 | 400 | 1.00 (0.00) | 1.00 (0.00) | 0.00 (0.00) | 0.00 (0.00) | 0.00 (0.00) |
| 15 | observed | 40 | 0.30 (0.19) | 0.30 (0.19) | 2.30 (12.12) | - | - |
| 16 | observed | 100 | 0.56 (0.29) | 0.55 (0.29) | 0.90 (7.23) | - | - |
| 17 | observed | 160 | 0.69 (0.30) | 0.68 (0.30) | 1.41 (6.37) | - | - |
| 18 | observed | 220 | 0.76 (0.29) | 0.75 (0.29) | 0.96 (5.39) | - | - |
| 19 | observed | 280 | 0.80 (0.28) | 0.80 (0.28) | 0.97 (5.18) | - | - |
| 20 | observed | 340 | 0.83 (0.27) | 0.83 (0.27) | 0.67 (4.54) | - | - |
| 21 | observed | 400 | 0.85 (0.26) | 0.85 (0.26) | 0.61 (4.26) | - | - |

**Simulation problem 6**

| | *Target effect size* | *Target sample size* | $\hat{\text{pwr}}^{\text{ups}}$ *mean (sd)* | $\hat{\text{pwr}}^{\text{cmp}}$ *mean (sd)* | $\text{PE}^{\text{ups,cmp}}$ *mean (sd)* | $\text{PE}^{\text{ups,true}}$ *mean (sd)* | $\text{PE}^{\text{cmp,true}}$ *mean (sd)* |
|---|---|---|---|---|---|---|---|

| | | | | | | | |
|---|---|---|---|---|---|---|---|
| 1 | 0.5 | 40 | 0.20 (0.02) | 0.20 (0.02) | -2.60 (9.79) | 1.32 (10.50) | 4.74 (12.50) |
| 2 | 0.5 | 100 | 0.40 (0.05) | 0.41 (0.05) | -0.66 (5.64) | 9.92 (14.36) | 10.88 (14.89) |
| 3 | 0.5 | 160 | 0.57 (0.07) | 0.57 (0.07) | -0.06 (4.18) | 8.69 (13.87) | 8.90 (14.21) |
| 4 | 0.5 | 220 | 0.70 (0.08) | 0.70 (0.08) | -0.40 (2.92) | 7.01 (12.07) | 7.50 (12.19) |
| 5 | 0.5 | 280 | 0.79 (0.07) | 0.79 (0.07) | -0.38 (2.44) | 6.70 (10.12) | 7.14 (10.13) |
| 6 | 0.5 | 340 | 0.85 (0.07) | 0.86 (0.07) | -0.32 (1.86) | 4.60 (8.20) | 4.95 (8.14) |
| 7 | 0.5 | 400 | 0.90 (0.06) | 0.90 (0.06) | -0.25 (1.59) | 2.36 (6.41) | 2.63 (6.34) |
| 8 | 1 | 40 | 0.53 (0.06) | 0.54 (0.08) | -0.42 (6.79) | 9.71 (13.00) | 10.86 (16.13) |
| 9 | 1 | 100 | 0.88 (0.06) | 0.89 (0.06) | -1.11 (1.63) | 2.48 (6.90) | 3.65 (6.96) |
| 10 | 1 | 160 | 0.97 (0.03) | 0.98 (0.02) | -0.52 (0.79) | 0.37 (2.79) | 0.89 (2.53) |
| 11 | 1 | 220 | 0.99 (0.01) | 0.99 (0.01) | -0.20 (0.40) | -0.14 (1.07) | 0.05 (0.88) |
| 12 | 1 | 280 | 1.00 (0.00) | 1.00 (0.00) | -0.07 (0.20) | -0.10 (0.42) | -0.03 (0.32) |
| 13 | 1 | 340 | 1.00 (0.00) | 1.00 (0.00) | -0.02 (0.10) | -0.03 (0.18) | -0.01 (0.12) |
| 14 | 1 | 400 | 1.00 (0.00) | 1.00 (0.00) | -0.01 (0.04) | -0.01 (0.06) | -0.01 (0.05) |
| 15 | observed | 40 | 0.24 (0.16) | 0.25 (0.17) | -1.23 (12.01) | - | - |
| 16 | observed | 100 | 0.42 (0.27) | 0.44 (0.28) | -2.04 (9.49) | - | - |
| 17 | observed | 160 | 0.54 (0.31) | 0.55 (0.31) | -1.07 (9.33) | - | - |
| 18 | observed | 220 | 0.61 (0.32) | 0.62 (0.32) | -0.61 (9.52) | - | - |
| 19 | observed | 280 | 0.66 (0.32) | 0.67 (0.32) | -0.34 (9.19) | - | - |
| 20 | observed | 340 | 0.70 (0.32) | 0.70 (0.32) | -0.07 (9.67) | - | - |
| 21 | observed | 400 | 0.72 (0.31) | 0.73 (0.32) | -0.12 (9.72) | - | - |

Table SM.2: Summary of power estimates and percentage errors (PEs) across simulation problems 7-9. The first and second table columns specify target proportion of dichotomous covariate $X_1 = 1$ and target sample size. Table columns third and fourth show aggregates (mean and standard deviation) of power estimates and PE across $R = 1000$ independent experiment repetitions: $\hat{\text{pwr}}^{\text{ups}}$ – upstrap power, $\text{PE}^{\text{cmp,true}}$ – percentage error between comparator and true power.

**Simulation problem 7**

| | *Target prop.* $X_1 = 1$ | *Target sample size* | $\hat{\text{pwr}}^{\text{ups}}$ *mean (sd)* | $\text{PE}^{\text{ups,true}}$ *mean (sd)* |
|---|---|---|---|---|
| 1 | 0.05 | 40 | 0.13 (0.05) | 5.28 (38.02) |
| 2 | 0.05 | 100 | 0.26 (0.07) | 3.59 (25.48) |
| 3 | 0.05 | 160 | 0.39 (0.07) | 2.21 (19.52) |
| 4 | 0.05 | 220 | 0.50 (0.08) | 1.28 (15.43) |
| 5 | 0.05 | 280 | 0.60 (0.08) | 1.71 (12.88) |
| 6 | 0.05 | 340 | 0.69 (0.07) | 1.05 (10.52) |
| 7 | 0.05 | 400 | 0.75 (0.07) | 0.65 (8.83) |
| 8 | 0.1 | 40 | 0.21 (0.05) | 8.03 (28.28) |
| 9 | 0.1 | 100 | 0.44 (0.07) | 3.26 (17.10) |
| 10 | 0.1 | 160 | 0.63 (0.07) | 2.47 (12.09) |
| 11 | 0.1 | 220 | 0.77 (0.06) | 0.54 (8.48) |
| 12 | 0.1 | 280 | 0.86 (0.05) | 0.70 (6.08) |
| 13 | 0.1 | 340 | 0.91 (0.04) | 0.42 (4.29) |
| 14 | 0.1 | 400 | 0.95 (0.03) | 0.27 (2.98) |
| 15 | 0.3 | 40 | 0.41 (0.06) | 2.02 (14.56) |
| 16 | 0.3 | 100 | 0.79 (0.06) | 0.89 (7.66) |
| 17 | 0.3 | 160 | 0.93 (0.03) | -0.22 (3.41) |
| 18 | 0.3 | 220 | 0.98 (0.01) | -0.31 (1.42) |
| 19 | 0.3 | 280 | 0.99 (0.01) | -0.04 (0.53) |
| 20 | 0.3 | 340 | 1.00 (0.00) | -0.02 (0.20) |
| 21 | 0.3 | 400 | 1.00 (0.00) | -0.02 (0.09) |
| 22 | 0.5 | 40 | 0.47 (0.06) | 3.46 (12.41) |
| 23 | 0.5 | 100 | 0.85 (0.05) | 0.88 (5.95) |
| 24 | 0.5 | 160 | 0.96 (0.02) | 0.28 (2.24) |
| 25 | 0.5 | 220 | 0.99 (0.01) | -0.23 (0.73) |
| 26 | 0.5 | 280 | 1.00 (0.00) | -0.06 (0.25) |
| 27 | 0.5 | 340 | 1.00 (0.00) | 0.01 (0.08) |
| 28 | 0.5 | 400 | 1.00 (0.00) | -0.01 (0.03) |

**Simulation problem 8**

| | *Target prop.* $X_1 = 1$ | *Target sample size* | $\hat{\text{pwr}}^{\text{ups}}$ *mean (sd)* | $\text{PE}^{\text{ups,true}}$ *mean (sd)* |
|---|---|---|---|---|
| 1 | 0.05 | 40 | 0.28 (0.06) | 9.48 (25.41) |
| 2 | 0.05 | 100 | 0.59 (0.08) | 5.01 (13.73) |
| 3 | 0.05 | 160 | 0.79 (0.06) | 1.82 (8.14) |
| 4 | 0.05 | 220 | 0.90 (0.04) | 0.18 (4.89) |
| 5 | 0.05 | 280 | 0.95 (0.03) | 0.15 (2.88) |

| | | | | |
|---|---|---|---|---|
| 6 | 0.05 | 340 | 0.98 (0.02) | 0.14 (1.63) |
| 7 | 0.05 | 400 | 0.99 (0.01) | -0.03 (0.91) |
| 8 | 0.1 | 40 | 0.46 (0.08) | 3.83 (17.00) |
| 9 | 0.1 | 100 | 0.85 (0.06) | 1.43 (6.78) |
| 10 | 0.1 | 160 | 0.96 (0.02) | -0.28 (2.53) |
| 11 | 0.1 | 220 | 0.99 (0.01) | -0.10 (0.92) |
| 12 | 0.1 | 280 | 1.00 (0.00) | -0.00 (0.32) |
| 13 | 0.1 | 340 | 1.00 (0.00) | -0.01 (0.11) |
| 14 | 0.1 | 400 | 1.00 (0.00) | 0.01 (0.04) |
| 15 | 0.3 | 40 | 0.79 (0.06) | 0.56 (7.70) |
| 16 | 0.3 | 100 | 0.99 (0.01) | -0.19 (0.67) |
| 17 | 0.3 | 160 | 1.00 (0.00) | -0.01 (0.05) |
| 18 | 0.3 | 220 | 1.00 (0.00) | -0.00 (0.01) |
| 19 | 0.3 | 280 | 1.00 (0.00) | 0.00 (0.00) |
| 20 | 0.3 | 340 | 1.00 (0.00) | 0.00 (0.00) |
| 21 | 0.3 | 400 | 1.00 (0.00) | 0.00 (0.00) |
| 22 | 0.5 | 40 | 0.85 (0.05) | 1.04 (6.08) |
| 23 | 0.5 | 100 | 1.00 (0.00) | 0.02 (0.30) |
| 24 | 0.5 | 160 | 1.00 (0.00) | -0.00 (0.02) |
| 25 | 0.5 | 220 | 1.00 (0.00) | 0.00 (0.00) |
| 26 | 0.5 | 280 | 1.00 (0.00) | 0.00 (0.00) |
| 27 | 0.5 | 340 | 1.00 (0.00) | 0.00 (0.00) |
| 28 | 0.5 | 400 | 1.00 (0.00) | 0.00 (0.00) |

**Simulation problem 9**

| | *Target prop.* $X_1 = 1$ | *Target sample size* | $\hat{\text{pwr}}^{\text{ups}}$ *mean (sd)* | $\text{PE}^{\text{ups,true}}$ *mean (sd)* |
|---|---|---|---|---|
| 1 | 0.05 | 40 | 0.23 (0.06) | 14.87 (29.98) |
| 2 | 0.05 | 100 | 0.48 (0.08) | 6.67 (17.59) |
| 3 | 0.05 | 160 | 0.68 (0.08) | 4.74 (12.12) |
| 4 | 0.05 | 220 | 0.80 (0.07) | 2.05 (8.37) |
| 5 | 0.05 | 280 | 0.88 (0.05) | 1.02 (5.94) |
| 6 | 0.05 | 340 | 0.93 (0.04) | 0.44 (4.22) |
| 7 | 0.05 | 400 | 0.96 (0.03) | -0.06 (2.97) |
| 8 | 0.1 | 40 | 0.37 (0.07) | 7.38 (20.45) |
| 9 | 0.1 | 100 | 0.74 (0.07) | 2.71 (9.94) |
| 10 | 0.1 | 160 | 0.90 (0.04) | 0.76 (4.92) |
| 11 | 0.1 | 220 | 0.97 (0.02) | -0.03 (2.29) |
| 12 | 0.1 | 280 | 0.99 (0.01) | -0.13 (1.13) |
| 13 | 0.1 | 340 | 1.00 (0.01) | -0.14 (0.58) |
| 14 | 0.1 | 400 | 1.00 (0.00) | -0.08 (0.30) |
| 15 | 0.3 | 40 | 0.67 (0.06) | 3.10 (9.92) |
| 16 | 0.3 | 100 | 0.97 (0.02) | 0.06 (1.89) |
| 17 | 0.3 | 160 | 1.00 (0.00) | -0.06 (0.28) |
| 18 | 0.3 | 220 | 1.00 (0.00) | -0.01 (0.04) |
| 19 | 0.3 | 280 | 1.00 (0.00) | -0.00 (0.01) |
| 20 | 0.3 | 340 | 1.00 (0.00) | -0.00 (0.00) |

| | | | | |
|---|---|---|---|---|
| 21 | 0.3 | 400 | 1.00 (0.00) | 0.00 (0.00) |
| 22 | 0.5 | 40 | 0.75 (0.06) | 1.36 (8.70) |
| 23 | 0.5 | 100 | 0.99 (0.01) | -0.09 (1.09) |
| 24 | 0.5 | 160 | 1.00 (0.00) | -0.03 (0.12) |
| 25 | 0.5 | 220 | 1.00 (0.00) | -0.00 (0.02) |
| 26 | 0.5 | 280 | 1.00 (0.00) | -0.00 (0.00) |
| 27 | 0.5 | 340 | 1.00 (0.00) | 0.00 (0.00) |
| 28 | 0.5 | 400 | 1.00 (0.00) | 0.00 (0.00) |

Table SM.3: Summary of power estimates and percentage errors (PEs) across simulation problems 10-12. The first, second ad third table columns specify observed sample size, target sample size, and assumed standard deviation of error term in data-generating model (the second largest error's standard deviation value only was selected for presentation in the table within each simulation problem). Table columns fourth and fifth show aggregates (mean and standard deviation) of power estimates and PE across $R = 1000$ independent experiment repetitions: $\hat{\text{pwr}}^{\text{ups}}$ – upstrap power, $\text{PE}^{\text{cmp,true}}$ – percentage error between comparator and true power.

**Simulation problem 10**

| | *Observed sample size* | *Target sample size* | *Error distr. std.* | $\hat{\text{pwr}}^{\text{ups}}$ *mean (sd)* | $\text{PE}^{\text{ups,true}}$ *mean (sd)* |
|---|---|---|---|---|---|
| 1 | 10 | 40 | 0.7 | 0.37 (0.17) | 43.77 (65.79) |
| 2 | 10 | 100 | 0.7 | 0.70 (0.19) | 25.43 (33.44) |
| 3 | 10 | 160 | 0.7 | 0.85 (0.14) | 10.08 (18.09) |
| 4 | 10 | 220 | 0.7 | 0.92 (0.10) | 3.45 (10.85) |
| 5 | 10 | 280 | 0.7 | 0.96 (0.07) | 1.06 (7.01) |
| 6 | 10 | 340 | 0.7 | 0.98 (0.05) | -0.07 (4.69) |
| 7 | 10 | 400 | 0.7 | 0.99 (0.03) | -0.34 (3.24) |
| 8 | 20 | 40 | 0.7 | 0.31 (0.09) | 20.48 (36.54) |
| 9 | 20 | 100 | 0.7 | 0.64 (0.14) | 14.94 (24.87) |
| 10 | 20 | 160 | 0.7 | 0.82 (0.11) | 6.26 (14.63) |
| 11 | 20 | 220 | 0.7 | 0.91 (0.08) | 2.16 (8.74) |
| 12 | 20 | 280 | 0.7 | 0.95 (0.05) | 0.75 (5.29) |
| 13 | 20 | 340 | 0.7 | 0.98 (0.03) | -0.03 (3.21) |
| 14 | 20 | 400 | 0.7 | 0.99 (0.02) | -0.19 (1.96) |
| 15 | 40 | 40 | 0.7 | 0.28 (0.06) | 9.24 (22.31) |
| 16 | 40 | 100 | 0.7 | 0.60 (0.10) | 7.80 (17.59) |
| 17 | 40 | 160 | 0.7 | 0.79 (0.09) | 2.89 (11.38) |
| 18 | 40 | 220 | 0.7 | 0.90 (0.06) | 0.74 (7.05) |
| 19 | 40 | 280 | 0.7 | 0.95 (0.04) | 0.20 (4.26) |
| 20 | 40 | 340 | 0.7 | 0.98 (0.02) | -0.21 (2.56) |
| 21 | 40 | 400 | 0.7 | 0.99 (0.02) | -0.20 (1.53) |
| 22 | 100 | 40 | 0.7 | 0.27 (0.04) | 4.23 (14.10) |
| 23 | 100 | 100 | 0.7 | 0.58 (0.06) | 3.99 (11.26) |
| 24 | 100 | 160 | 0.7 | 0.78 (0.06) | 1.02 (7.58) |
| 25 | 100 | 220 | 0.7 | 0.89 (0.04) | -0.05 (4.87) |
| 26 | 100 | 280 | 0.7 | 0.95 (0.03) | -0.00 (2.94) |
| 27 | 100 | 340 | 0.7 | 0.98 (0.02) | -0.14 (1.68) |
| 28 | 100 | 400 | 0.7 | 0.99 (0.01) | -0.09 (0.93) |

**Simulation problem 11**

| | *Observed sample size* | *Target sample size* | *Error distr. std.* | $\hat{\text{pwr}}^{\text{ups}}$ *mean (sd)* | $\text{PE}^{\text{ups,true}}$ *mean (sd)* |
|---|---|---|---|---|---|
| 1 | 10 | 40 | 1 | 0.45 (0.23) | 41.50 (72.51) |
| 2 | 10 | 100 | 1 | 0.79 (0.20) | 16.27 (29.51) |
| 3 | 10 | 160 | 1 | 0.91 (0.13) | 4.57 (15.27) |

| | | | | | |
|---|---|---|---|---|---|
| 4 | 10 | 220 | 1 | 0.96 (0.09) | 0.78 (9.51) |
| 5 | 10 | 280 | 1 | 0.98 (0.07) | -0.74 (6.64) |
| 6 | 10 | 340 | 1 | 0.99 (0.05) | -0.72 (5.03) |
| 7 | 10 | 400 | 1 | 0.99 (0.04) | -0.67 (4.05) |
| 8 | 20 | 40 | 1 | 0.38 (0.12) | 17.00 (37.76) |
| 9 | 20 | 100 | 1 | 0.75 (0.14) | 9.66 (20.67) |
| 10 | 20 | 160 | 1 | 0.90 (0.09) | 3.25 (10.75) |
| 11 | 20 | 220 | 1 | 0.96 (0.06) | 0.86 (5.95) |
| 12 | 20 | 280 | 1 | 0.98 (0.03) | -0.30 (3.39) |
| 13 | 20 | 340 | 1 | 0.99 (0.02) | -0.27 (2.10) |
| 14 | 20 | 400 | 1 | 1.00 (0.01) | -0.28 (1.34) |
| 15 | 40 | 40 | 1 | 0.35 (0.08) | 7.97 (23.83) |
| 16 | 40 | 100 | 1 | 0.72 (0.10) | 5.38 (15.05) |
| 17 | 40 | 160 | 1 | 0.89 (0.07) | 2.07 (8.02) |
| 18 | 40 | 220 | 1 | 0.96 (0.04) | 0.71 (4.10) |
| 19 | 40 | 280 | 1 | 0.98 (0.02) | -0.13 (2.01) |
| 20 | 40 | 340 | 1 | 0.99 (0.01) | -0.06 (0.98) |
| 21 | 40 | 400 | 1 | 1.00 (0.00) | -0.11 (0.49) |
| 22 | 100 | 40 | 1 | 0.33 (0.04) | 2.97 (13.21) |
| 23 | 100 | 100 | 1 | 0.70 (0.06) | 2.49 (9.33) |
| 24 | 100 | 160 | 1 | 0.88 (0.05) | 1.14 (5.17) |
| 25 | 100 | 220 | 1 | 0.96 (0.02) | 0.69 (2.59) |
| 26 | 100 | 280 | 1 | 0.98 (0.01) | -0.04 (1.25) |
| 27 | 100 | 340 | 1 | 0.99 (0.01) | 0.03 (0.56) |
| 28 | 100 | 400 | 1 | 1.00 (0.00) | -0.05 (0.24) |

**Simulation problem 12**

| | *Observed sample size* | *Target sample size* | *Error distr. std.* | $\hat{\text{pwr}}^{\text{ups}}$ *mean (sd)* | $\text{PE}^{\text{ups,true}}$ *mean (sd)* |
|---|---|---|---|---|---|
| 1 | 10 | 40 | 1 | 0.35 (0.19) | 39.31 (76.73) |
| 2 | 10 | 100 | 1 | 0.69 (0.23) | 23.57 (41.96) |
| 3 | 10 | 160 | 1 | 0.84 (0.19) | 9.22 (24.24) |
| 4 | 10 | 220 | 1 | 0.91 (0.14) | 2.76 (15.88) |
| 5 | 10 | 280 | 1 | 0.95 (0.11) | 0.03 (11.21) |
| 6 | 10 | 340 | 1 | 0.97 (0.08) | -0.69 (8.36) |
| 7 | 10 | 400 | 1 | 0.98 (0.06) | -1.04 (6.45) |
| 8 | 20 | 40 | 1 | 0.30 (0.10) | 18.19 (38.85) |
| 9 | 20 | 100 | 1 | 0.64 (0.15) | 13.91 (27.10) |
| 10 | 20 | 160 | 1 | 0.82 (0.13) | 6.62 (16.54) |
| 11 | 20 | 220 | 1 | 0.91 (0.09) | 2.62 (10.27) |
| 12 | 20 | 280 | 1 | 0.95 (0.06) | 0.68 (6.55) |
| 13 | 20 | 340 | 1 | 0.97 (0.04) | 0.12 (4.25) |
| 14 | 20 | 400 | 1 | 0.99 (0.03) | -0.25 (2.78) |
| 15 | 40 | 40 | 1 | 0.28 (0.06) | 8.84 (23.92) |
| 16 | 40 | 100 | 1 | 0.60 (0.10) | 7.35 (18.73) |
| 17 | 40 | 160 | 1 | 0.80 (0.09) | 3.73 (12.16) |
| 18 | 40 | 220 | 1 | 0.90 (0.07) | 1.69 (7.61) |

| | | | | | |
|---|---|---|---|---|---|
| 19 | 40 | 280 | 1 | 0.95 (0.04) | 0.62 (4.67) |
| 20 | 40 | 340 | 1 | 0.98 (0.03) | 0.31 (2.90) |
| 21 | 40 | 400 | 1 | 0.99 (0.02) | -0.02 (1.80) |
| 22 | 100 | 40 | 1 | 0.26 (0.04) | 3.46 (13.86) |
| 23 | 100 | 100 | 1 | 0.58 (0.06) | 3.44 (11.21) |
| 24 | 100 | 160 | 1 | 0.78 (0.06) | 1.80 (7.70) |
| 25 | 100 | 220 | 1 | 0.89 (0.04) | 0.98 (4.90) |
| 26 | 100 | 280 | 1 | 0.95 (0.03) | 0.45 (2.93) |
| 27 | 100 | 340 | 1 | 0.98 (0.02) | 0.41 (1.68) |
| 28 | 100 | 400 | 1 | 0.99 (0.01) | 0.11 (0.94) |
